# Linking place and view: Organizing space through saccades and fixations between primate posterior parietal cortex and hippocampus

**DOI:** 10.1101/2023.11.06.565451

**Authors:** Marie Vericel, Pierre Baraduc, Jean René Duhamel, Sylvia Wirth

## Abstract

Humans primarily rely on vision to explore and guide actions in spatial environments. The parietal cortex is thought to withhold a unified representation of the visual space allowing to direct saccades to salient cues, while the hippocampus provides a memory-based cognitive place map of the environment. Understanding how these two representations interact during navigation is a key question. To probe the link between view and place, we compared neural activity in the posterior parietal cortex and hippocampus of macaques navigating in a virtual maze. When analyzed as a function of the animal’s position in the virtual environment, more neurons in the parietal cortex displayed spatial selectivity compared to the hippocampus. We hypothesized that such modulation by self-position in the parietal cortex might stem from processing visual cues of the environment through exploratory saccades and fixations. However, we established that position-selectivity was not solely correlated with simple oculomotor dynamics. Rather, spatial selectivities in the PPC and the HPC originated from cells driven by direct fixations of maze paths or landmarks. However, while a substantial proportion of PPC and HPC cells displayed selectivity towards landmarks’ features, such as their side of appearance or their identity, we also revealed different task-related maze segmentation between regions. Indeed, when animal gazed at paths, activity in parietal cortex revealed anticipation of reward while that of the hippocampus suggested reward outcome processing. On the other hand, when animals gazed at a landmark already present in the field of view, parietal activity tended to occur close to intersections, while that of hippocampus was more spatially distributed. Finally, at the population level, neurons in both regions anticipated landmarks before they appeared in the field of view, suggesting a shared knowledge of the spatial layout and a collective active role in memory-guided visual exploration across regions. Taken together, these findings shed light on the neural processes that link place and view, through action- and memory-driven exploration of objects in space.

## Introduction

While we navigate skillfully from our desks into virtual space, this competence is not a human prerogative. Non-human primates and rodents can find their way in virtual environments too, avoiding obstacles and planning trajectories to reach goals (Acharya et al. 2016; Chen et al. 2013; Corrigan et al. 2017; Cushman et al. 2013; Gulli et al. 2020; Harvey et al. 2012; Hori et al. 2005; Sato et al. 2004; Taillade et al. 2019; Wirth et al. 2017). Then, how can a sense of space arise from visual-only stimulation? Optic flow suffices to create an illusion of self-motion and to provide heading informing on travel direction (Li and Warren 2000). Further, visual processing of a scene alone allows to identify relevant cues for navigation, such as objects, landmarks, paths or borders, providing goals to reach, and framing them in self-based or other-objects-based spatial coordinates (Arleo and Rondi-Reig 2007; Bicanski and Burgess 2020; Ekstrom 2015; Wolbers et al. 2008). In comparison with the place-selective cells typically described in rodents hippocampus (HPC; O’Keefe and Burgess 1996; O’Keefe and Dostrovsky 1971; O’Keefe and Nadel 1979), it has been shown that primate HPC neurons are active when the animal looks at specific landmarks and views of the environment, in a task-driven manner during virtual navigation (Corrigan et al. 2023; Gulli et al. 2020; Wirth et al. 2017) or random foraging in real-world (Mao et al. 2021), suggesting a role in the encoding of a map of the visual space. However, its processing by saccades and fixations is known to arise from neural computations performed in the posterior parietal cortex (PPC), notably in the intraparietal areas (Bremmer 2005; Cohen and Andersen 2002; Colby and Duhamel 1991; Kravitz et al. 2011). In the lateral and the ventral intraparietal areas (LIP and VIP), these computations aim at tracking self- and objects motions, and optic-flow direction and velocity (Bremmer et al. 2002a; Chen et al. 2011; Sunkara et al. 2016), the direction of moving stimuli (Bremmer et al. 2002b; Duhamel et al. 1998; Fanini and Assad 2009), or their saliency (Ben Hamed 2002; Goldberg et al. 2006; Zhang et al. 2014). Further, they encode spatial coordinates of visual stimuli in a continuum between eye- and head-centered reference frames (Avillac et al. 2005; Cohen and Andersen 2002; Duhamel et al. 1997; Mullette-Gillman et al. 2005). Finally, LIP also supports a continuous and stable visual space taking into account the programming of ocular saccades during visual exploration (Bremmer et al. 2016; Duhamel et al. 1992; Fanini and Assad 2009; Hagan et al. 2012; Heiser and Colby 2006; Platt and Glimcher 1999; Steenrod et al. 2013; Zhou et al. 2018). In sum, the studies suggest that parietal cortex, along with hippocampal regions, may greatly be recruited during navigational tasks (Arbib 1997; Baumann and Mattingley 2014; Bremmer 2005). However, while rodents PPC neurons were shown to display route-selective activities (Nitz 2006; Shelley et al. 2022) or heading angle (Krumin et al. 2018), there have been no studies that characterized activity in intraparietal regions during landmark-based navigation in the non-human primate.

Anatomically, parietal inputs to hippocampus take several polysynaptic pathways via the retrosplenial, posterior cingulate, or the parahippocampal cortices before reaching entorhinal cortex or subiculum (Insausti et al. 1987; Kobayashi and Amaral 2003, 2007; Van Hoesen 1982). Further, the hippocampal formation projects back to parietal regions via the reverse path (Kobayashi and Amaral 2003, 2007) or directly (Van Hoesen 1982). These data, together with lesioning experiments in rodents (Ammassari-Teule et al. 1998; Rogers and Kesner 2007; Save et al. 2005; Thinus-Blanc et al. 1996), provide more ground to support an interplay between hippocampus and PPC for navigation (Burgess et al. 1997; Save and Poucet 2000).

To understand the nature of neural activity in PPC during a goal-directed, virtual-reality task, we compared the activity of cells of LIP and VIP with that of hippocampal ones, as a function of virtual position, and as a function of saccades and fixations to landmarks. We hypothesized that a “place” representation in PPC would derive from the processing of relevant salient cues of the visual space, and that this representation would complement the role of hippocampus in cues identification and environment mapping. Our results show how parietal cortex and hippocampus displayed position-related activities resulting from a dynamic processing of visual cues at both different and shared strategic task moments.

## Results

### Parietal cells display position selectivity as strongly as in the hippocampus

We analysed the activity of 111 cells recorded by laminar U-probes (Plexon®) in a continuum between the lateral and ventral banks of the intraparietal sulcus (PPC cells; Figure S1A-B) and 142 cells recorded throughout the whole hippocampus in the right hemisphere (HPC cells; the anatomical coordinates of the recording sites can be found in (Wirth et al. 2017), while macaque monkeys navigated a virtual 5 arms star-maze to obtain a hidden reward. Animals learned to locate the reward based on its position relative to five salient landmarks (see Figure 1A and Wirth et al., 2017). In all figures, we adopted the convention to orient the maze with the reward at North. During the session, animals started each trial by moving from the beginning of one of four inbound paths towards the centre of the maze (Figure 1A, steps 1-2, purple paths). Then they chose to enter another path with the joystick and learned via trial and error which path led to the reward (steps 3-4, red path). Following the reward delivery, they were passively allocated to a new start point through the “outbound” paths (step 5, dotted lines). The first-person view was uninterrupted between the trials (see bottom panel of Figure 1A). Following an initial shaping (see Methods) aimed to train the animals on the task rules, the landmarks identities were changed every day and monkeys quickly solved the task within each session, reaching 90% of correct trial after about 20 trials (Wirth et al. 2017).

**Figure 1.**
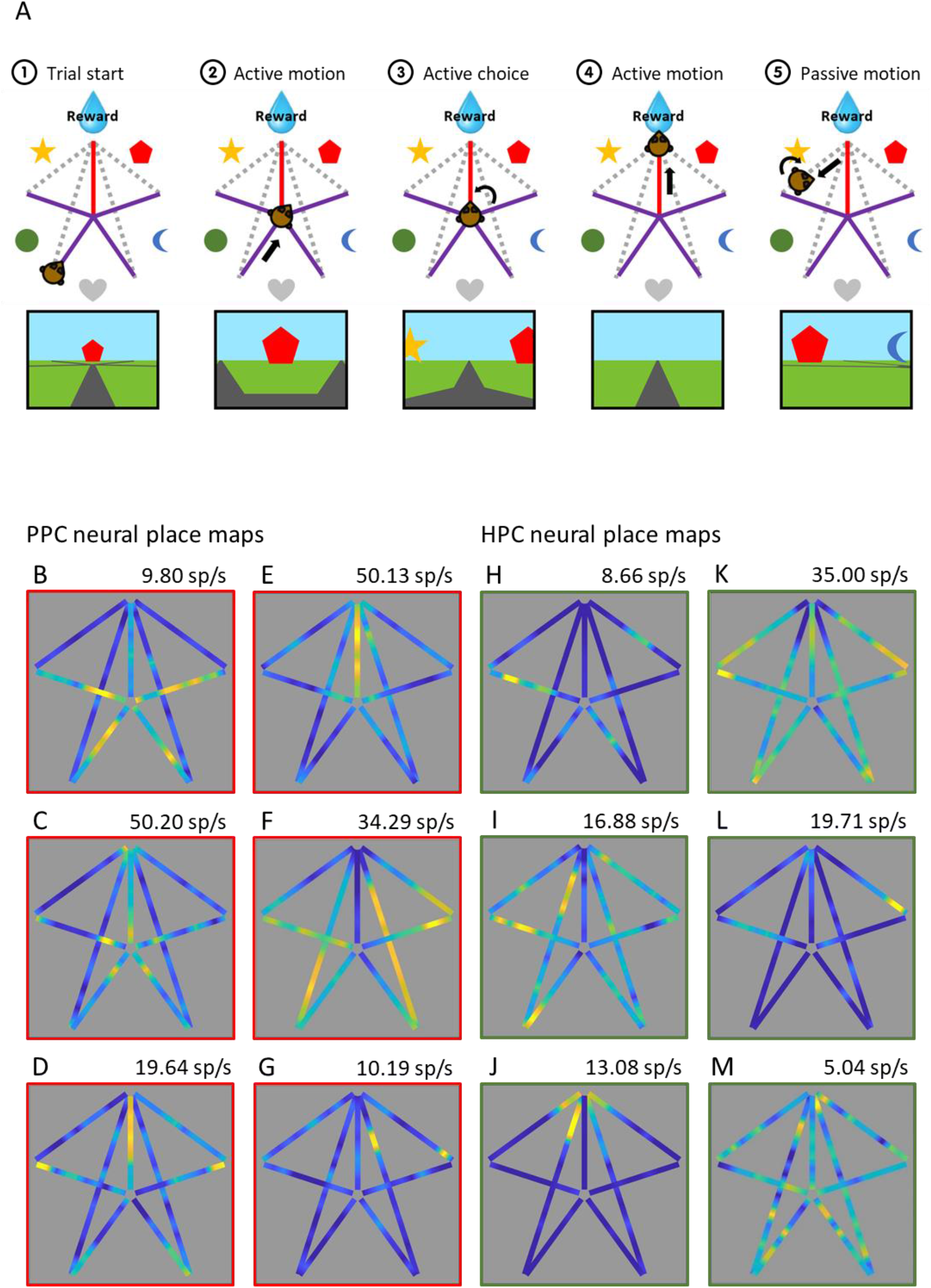
Recording sites and spatial selectivity in parietal and hippocampal neurons. A: Main steps of the navigation task performed by animals during the recordings, with their corresponding points of view below: 1) the trial began from the extremity of 1 of the 4 inbound paths of the star-maze (purple); 2) the animal pushed the joystick to move the camera and reach the maze center; 3) in the center, they pushed the joystick toward left or right as many time as they wanted, and then chose an exit path by pushing it forward; 4) if animals chose the correct path (red), they obtained a reward in the form of a few millimeters of water; 5) next, the animal was passively moved to the next starting position through the outbound paths (grey dashed lines), by an automatic movement of rotation and translation of the camera, and the next trial began. During the whole session, the first person view of the scene (bottom panels schematics of the view) was uninterrupted between trials. Five landmarks placed between the inbound paths were the only visual cues allowing orientation into the environment, since nothing was directly indicating the position of the reward. The task was presented on a large 152x114cm screen using a projector, and in stereopsis via a pair of shutter glasses synchronized with the projector. B-G: Heat maps of the firing rate of single unit examples of PPC neurons, as a function of monkey’s position into the star-maze (i.e. “neural place maps”). The color axis represents the firing rate (maximal firing rate, in spikes/s, indicated on the top left of each map). All cells had a significant position information content (IC). For a better visualization, data are displayed from the 5th to the 99th percentiles. H-M: Same conventions as for B-G, for HPC example cells. All cells had a significant position information content (IC).

We first examined whether parietal or hippocampal cells were modulated by animal’s virtual position, i.e. the camera’s location within the environment, independent from its orientation (see “neural place maps” on Figure 1B-M and S1C-J, with PPC maps outlined in red, and HPC in green; see Methods). Not only were parietal cells more active than hippocampal ones (mean firing rate of PPC=6.19±1.89 spikes/sec, µFR_HPC_=2.97±0.75 sp/s; Wilcoxon rank sum: Z=4.04, p=5.27×10^-5^), but more PPC cells displayed activity significantly modulated by position (91/111, 82.0%; defined as information content above that obtained from permutations, see Methods) than in the hippocampus (64/142, 45.1%; homogeneity Pearson’s Chi-squared with Yates’ correction: X²_1df_ =34.23, p=4.90×10^-9^; see Figures 1F-Q for examples of cells with significant modulation by position, Figures S1C-D for examples of cells with activity not significantly modulated by position). Comparison of various measures of selectivity such as the Information Content (IC; µIC_PPC_=0.32±0.059, µIC_HPC_=0.38±0.11; Wilcoxon rank sum: Z=0.49, p=0.62), sparsity indexes (see Methods; µS_PPC_=0.31±0.038, µS_HPC_=0.33±0.065; Wilcoxon rank sum: Z=0.46, p=0.65), or depth of tuning indexes (see Methods; µDT_PPC_=0.91±0.028, µDT_HPC_=0.89±0.043; Wilcoxon rank sum: Z=0.092, p=0.93) of the position-selective cells showed that parietal neurons were as selective to virtual position as hippocampal cells. In both areas, cells often expressed multiple spatial response fields (see Methods), but their number didn’t differ (µN_PPC_=2.96±0.26 RFs, µN_HPC_=2.87±0.28 RFs; Wilcoxon rank sum: Z=0.45, p=0.66). However, apart from a greater spatial response in the rewarded path, that was highly represented across the parietal population (see example in Figure 1E), visual inspection suggested that PPC cells expressed response fields in similar positions across paths: cells fired for instance at entry positions for all 4 inbound paths, as in example Figure 1D, or in the end of the inbound paths as in Figure 1B and C (see more symmetric responses in Figure S1E-F). To assess this apparent symmetry for all the position-modulated cells, we calculated separately the correlation coefficients (r) for activity in all inbound and all outbound paths. These correlation coefficients were modestly but significantly higher in the PPC (µr=0.15±0.034) than in the HPC (µr=0.099±0.041; Wilcoxon rank sum: Z=2.27, p=0.024). We also found a substantial number of cells that displayed asymmetric activity. Indeed, dividing the star-maze along the vertical axis centered on the rewarded path, a substantial fraction of the position-selective PPC (36/91, 39.6%) and HPC (26/64, 40.6%) cells showed greater activity in one side than the other for inbound paths, outbound or both (Wilcoxon rank sum: p≤0.05, see Figure S1G-J). However, at the population level, and for either the inbound paths, the outbound or the whole maze, we observed no preferences for one side over the other, neither in the PPC (Wilcoxon matched pairs signed-rank: Z_inbound_=1.24, p_inbound_=0.22; Z_outbound_=0.063, p_outbound_=0.95; Z_whole_=0.25, p_whole_=0.81; see Methods) nor in the HPC (Z_inbound_=-0.31, p_inbound_=0.75; Z_outbound_=0.094, p_outbound_=0.93; Z_whole_=-0.73, p_whole_=0.47). This suggests that the preferences observed at the single unit level were overall counterbalanced and thus that no relation between asymmetry and the recorded hemisphere could be found.

In sum, parietal neurons displayed strong modulation as a function of virtual position, and the pattern of position-related activity in individual examples suggested that PPC neurons were more recruited for specific task-states, regardless of the path identity, than neurons in the hippocampus. We then sought to determine whether these spatial selectivities were attributed to bottom-up visual information or to the top-down systematic exploratory behavior of the animals, with respect to the visual maze elements.

### Position selectivity is not exclusively driven by saccade-related activity

A position in virtual reality is solely provided by specific position-dependent visual information, which could drive specific behaviors, such as explorative saccades and fixations. Indeed, saccades occurred at an average rate of 2.44±0.11 sacc/s, with peaks at 6.25±0.26 sacc/s. Moreover, the individual (see Figure S2A-L), and averaged (Figure 2A) heat maps of the saccades’ frequencies as a function of monkey’s position (saccades maps; see Methods) show that they were mostly performed 1) on inbound paths before the center of the maze, 2) in the rewarded path and 3) at the end of the outbound paths, thereby reflecting gazing at landmarks and reward anticipation. This inhomogeneity was supported by a significant difference in saccade frequency (see Figure 2B, top panel; Kruskal-Wallis: X²_3df_ =41.70, p=4.64×10^-9^) and duration (see Figure 2B, bottom panel; Kruskal-Wallis: X²_3df_=18.8, p=3.01×10^-4^) between the four portions of the maze. Therefore, a factor potentially accounting for position neural selectivity lies in i) the link between the saccade-driven neural activity, ii) the explorative function of the saccades, and iii) the maze positions in which those saccades were performed due to the presence of key visual information.

**Figure 2.**
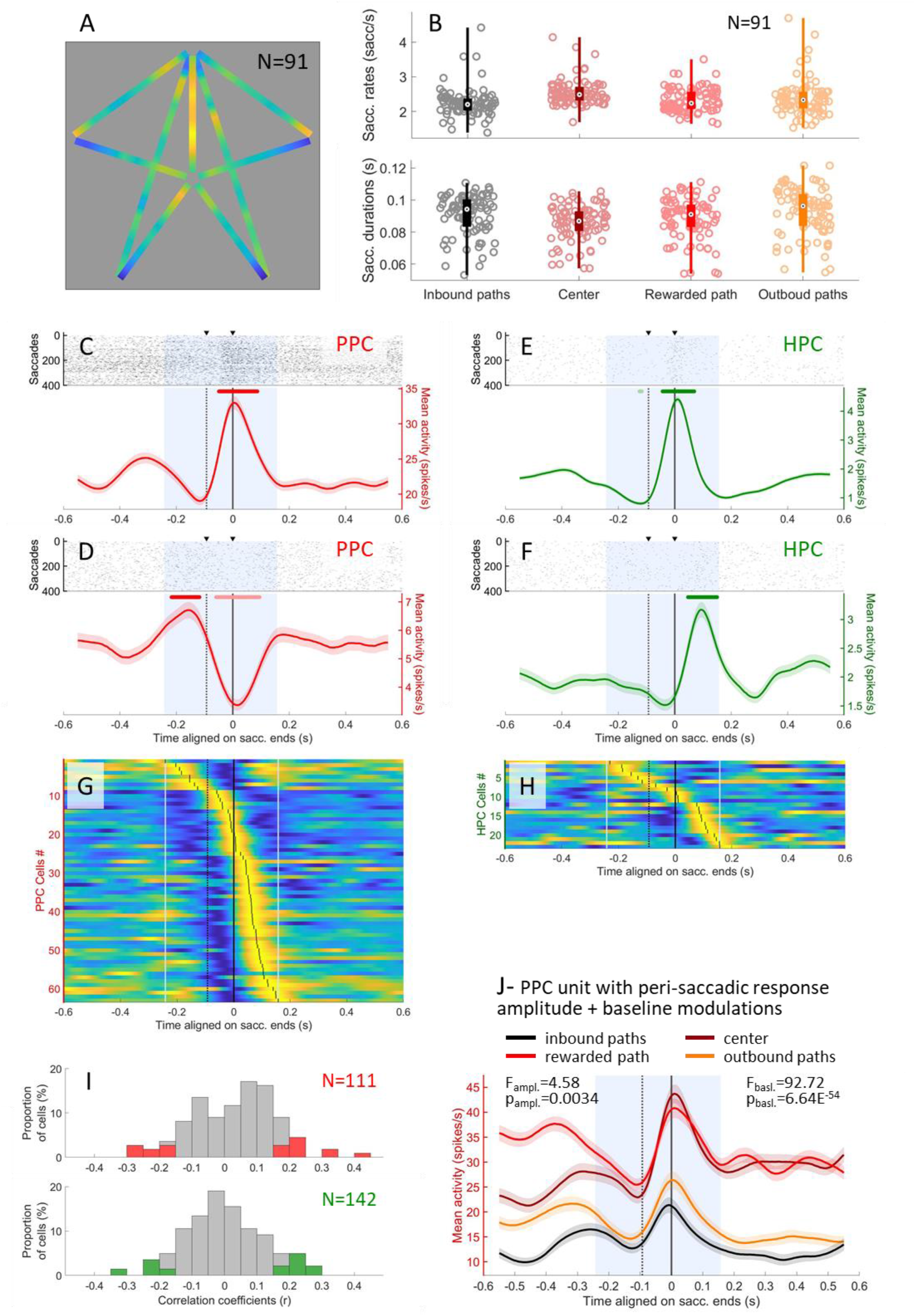
Saccade-related activities in posterior parietal and hippocampal neurons. A: Averaged of the normalized position heat map for behavioral saccades, for all the sessions recorded (N=91). The color axis represents the frequency of saccades (saccades/sec) as a function of the animal’s position in the maze. B: Scatter plots of the saccade rates (top) and durations (bottom) of each session (N=91), depending on the monkey’s position in the maze. The saccades rates, and durations were significantly different between the 4 maze segments (rates: Kruskal-Wallis: X²_3df_ =41.70, p=4.64×10^-9^; durations: X²_3df_ =18.8, p=3.01×10^-4^). C-D: Raster plot (top) and average activity (bottom, in red) of PPC neurons showing a peri-saccadic response when aligned on saccades ends (black solid line). The black dotted line represents the average time of saccades starts (-93ms). The standard errors of the mean are indicated in light color. The analysis window (-150 to +250ms relative to saccade start) is represented in a blue shaded area. The line above indicates times for which the response activity was significantly higher (dark; for A: from -50 to 87 ms; for B: from -218 to -118 ms) or lower (light; for B: from -60 to 95 ms) than baseline. E-F: Same as C-D, for hippocampus example cells. The activity was significantly higher than baseline from -44 to 69 ms relative to saccades ends for E, and from 46 to 150 ms for F, and lower than baseline from -126 to -117 ms for E. G: Population maps of the normalized activity of the PPC saccade-responsive cells (N=63), sorted by temporal order of their peak in the response window (-150 to +250ms relative to saccades starts, light blue dashed lines). Saccades ends are indicated by a black solid line, and starts by a black dotted line. H: Same as G, for the hippocampal saccade-responsive cells (N=23). I: Distribution of the Pearson correlation coefficients (r) of the correlation between the saccade frequency maps and the neural place map of each PPC (N=111, on top) and HPC (N=142, on bottom) neuron. The colored bars indicate the distribution of the r of significant correlations (determined by a permutation test, p≤0.05). 18/111 (16.2%) of the PPC cells (in red), and 22/142 (15.5%) of the HPC ones (in green) had their neural map significantly correlated with the saccades map. J: Averaged activity of an example PPC cell (same as in C), aligned on saccades ends (black solid line). The black dotted line represents the average time of saccades starts. The standard errors of the mean are indicated in light colors. The peri-saccadic activity is separated by segments in which the saccades are made (inbound paths in black, center in burgundy, rewarding path in red, outbound paths in orange). The cell displayed a significant difference of response peak amplitude (1-way-ANOVA: F_3df_=4.58, p=0.0034) as well as a significant difference in peri-saccadic response rate (1-way-ANOVA: F_3df_=92.72, p=6.64×10^-54^).

In line with the known role of PPC in saccade planning and fixations (Duhamel et al. 1992; Fanini and Assad 2009; Hagan et al. 2012; Heiser and Colby 2006; Platt and Glimcher 1999; Steenrod et al. 2013; Zhou et al. 2018), we found that 63/111 cells (56.8%) displayed significant peri-saccadic activity when aligned on saccades ends (see Methods), as shown in example Figures 2C-D. Surprisingly, hippocampal cells also showed peri-saccadic activities (23/138, 16.7%; Figure 2E-F), yet in a lower proportion than in the PPC (homogeneity Pearson’s Chi-squared with Yates’ correction: X²_1df_ =41.98, p=9.24×10^-11^). The averaged saccade frequency map did not differ between sessions in which responsive cells and the non-responsive cells were identified (Pearson correlation: r=0.95, p=1.40×10^-58^), indicating that the difference in behavior was not driving the differences in responsive and non-responsive cells’ activity. The peri-saccadic rate modulation could be positive (21/63, 33.3% of PPC responsive cells, 14/23, 60.9% of HPC ones; as shown in Figure 2C and F), negative (16/63, 25.4% in PPC, 8/23, 34.8% in HPC), or mixed (26/63, 41.3% in PPC, 1/23, 4.4% in HPC; Figure 2D and E). The population maps showed the same peri-saccadic temporal dynamics in PPC and HPC (see Figure 2G and H, respectively, with peak times indicated by black vertical ticks; Kolmogorov-Smirnov test on the distribution of the peaks: D=0.24, p=0.25, or latencies), with groups of pre-(N_PPC_=8/63, 12.7%; N_HPC_=6/23, 26.1%), trans-(N_PPC_=11/63, 17.5%; N_HPC_=2/23, 8.7%), and, for the majority, post-(N_PPC_=44/63, 69.8%; N_HPC_=15/23, 65.2%) saccade-active neurons (Figure S2M). In PPC, we found no link between the peak response time and the anatomical dorso-ventral coordinates of the neurons within the IPS (see Methods; Figure S2N; Pearson correlation: r_monkey K_=0.22, p_monkey K_=0.16; r_monkey S_=-0.097, p_monkey S_=0.68), and were thus unable to clearly separate LIP from the VIP neurons.

Next, to test whether virtual position neural maps were linked to saccade-frequency maps, we computed the correlations between these, calculated for individual cells. Figure S2A-F and G-L show the saccades maps of the cells presented in Figure 1B-G and H-M, respectively. We found that 19/111 (17.1%) of PPC cells and 22/142 (15.5%) of HPC ones showed a significant positive or negative correlation between their neural and saccades maps (Figure 2I; permutation test, p≤0.05; see Methods). Overall, the absolute values of correlation coefficients were significantly higher in PPC (µ|r|=0.11±0.015) than in HPC (µ|r|=0.095±0.012; tailed-Wilcoxon rank sum: Z=1.65, p=0.049), showing that parietal cells tended to be more positively or negatively saccade-responsive than HPC ones. In sum, the position neuronal map captures saccade-related activity, but only for a relatively small fraction of the cells.

An alternate possibility is that the context in which saccades and fixations were made actually impacted neuronal activity. To test this, for each cell, we compared the peri-saccadic activity (see Methods) across the four segments of the maze, i.e. the inbound paths, the center, the rewarded path and the outbounds. We found significant differences between activity depending on context. First, for a minority of cells, there was a modulation of the amplitude of the saccadic response *per se*, in PPC only (18/63, 28.6% of responsive cells; one-way-ANOVA: 1df, p≤0.05; example units in Figures 2J and S2O). This effect could stem from the difference in saccades durations observed between the four segments of the maze, described above (Figure 2B, bottom panel). However, we found that only 5 out of 18 neurons exhibited the highest firing rate in the segments in which saccade durations were also the highest, which is around the chance level (4.5). Together, those results imply that the kinematics of eye movements would not solely drive their activities. Second, for the majority of the cells, there was a modulation of the response baseline level depending on position in the maze, without necessarily a modulation of saccade-evoked response amplitude per se. Indeed, 55/63 (87.3%) of PPC and 17/23 (73.9%) of HPC saccade-responsive cells displayed a significant response rate difference across segments of the maze (Figures 2J, S2O and S2P; one-way-ANOVA, p≤0.05). At the population level, the preferred positions of those cells were uniformly distributed between the 4 maze segments (Figure S2Q), with 13/55 (23.6%) PPC cells displaying higher peri-saccadic activity in inbound paths, 18 (32.7%) in the center, 13 (23.6%) in the rewarded path, and 11 (20.8%) in the outbound paths (Chi-squared goodness-of-fit on a uniform theoretical distribution: X²_3df_=1.95, p=0.58). In hippocampus, 4/17 (23.5%) cells were more active in inbound paths, 2 (11.8%) in center, 5 (29.4%) in rewarded path and 6 (35.3%) in outbound paths (insufficient sample size to perform Chi-squared goodness-of-fit test). Thus, the PPC cells responded differently, depending on the ongoing task context, but no general attentional or motivational processes seemed to drive the whole neuronal population. Rather, each cell specifically responds to a specific sensori-motor context.

Overall, these results show that the heterogeneity in spatial activity patterns observed for single-cell neural place maps (Figure 1B-M and S1C-J) could only be partially imputed to pure oculomotor dynamics, but were also modulated by task context. Among the main factors likely varying with task context, we next analysed neurons’ activity as a function of directed gaze within the visual scene.

### A spectrum of responses to visual layout gives rise to different spatial selectivities across regions

As the visual landscape making up the virtual environment was simple, we constructed a continuous representation of the fields of view (FOV), allowing plotting the gaze on this panoramic scene (Figure 3A, left and middle panels), comprising the 4 landmarks neighbouring each of the paths (the 5^th^ one is usually not visible in correct trials, see Methods). The calculation of the point of gaze took into account the camera’s position and orientation, combined with the eye positions in the FOV. The point of gaze was then projected on a cylindrical wall at the landmark’s position, which we linearized into a panorama centered on the goal (Figure 3A). To assess which visual cues animals attended in the virtual environment, we quantified landmark or path fixations (see Methods). Animals fixated a landmark 0.18±0.0066 times per second (i.e. a landmark was foveated every 5.49 s). By comparison, animals fixated a path – excluding the rewarded path – 0.15±0.0096 times/s (i.e. a foveation every 6.84 s), which is significantly less than landmarks (Wilcoxon matched pairs signed-rank: Z=5.25, p=1.52×10^-7^). That animals preferred to direct their gaze to landmarks is in line with the hypothesis that these are more informative than paths.

**Figure 3.**
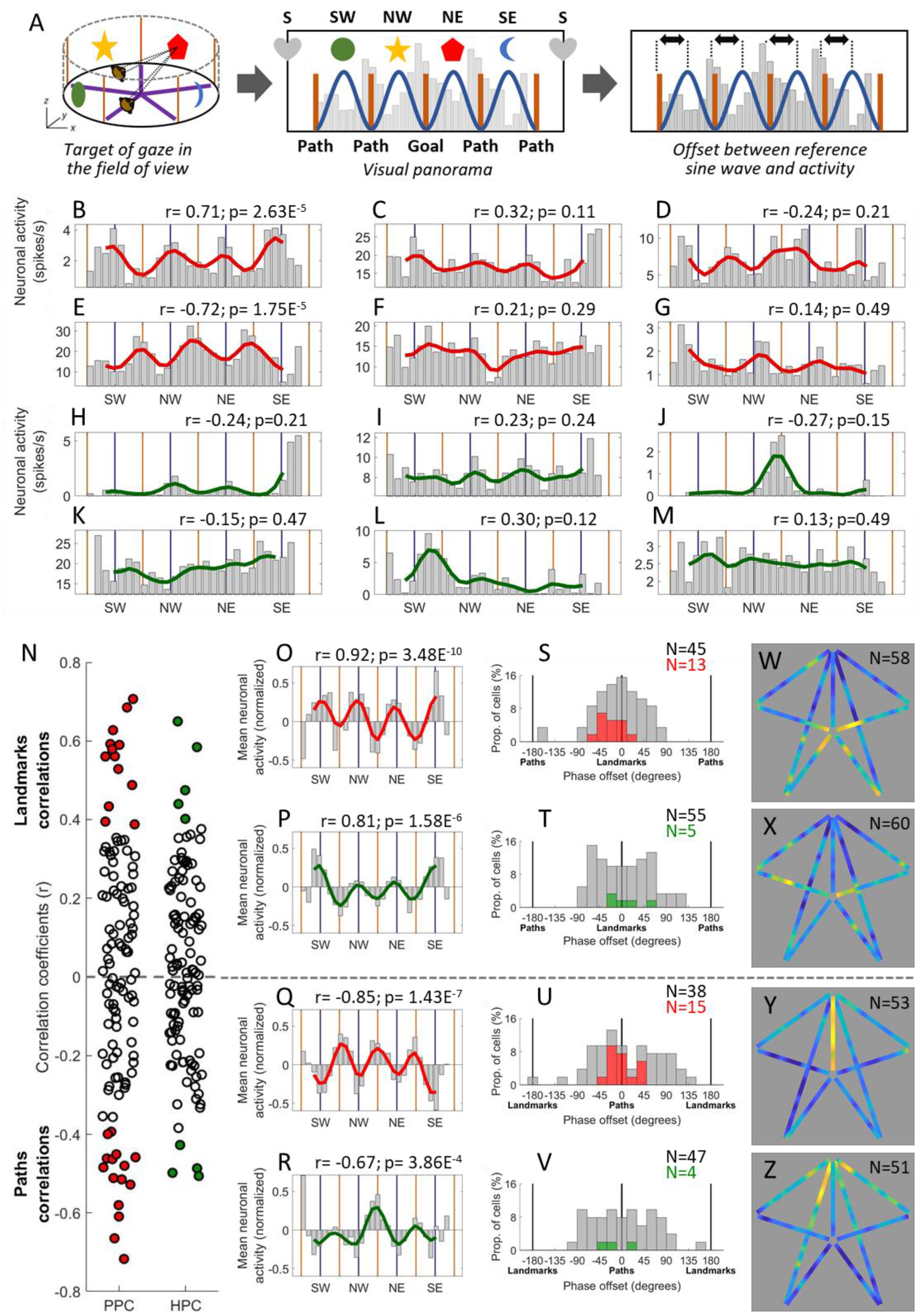
Activity as a function of the point of gaze. A: method used to analyse neuron’s firing rates as a function of monkey’s gaze position in the virtual environment: the point of gaze (left panel) in the visual panorama was computed by taking into account 1) monkey’s virtual facing location as a function of his position and head direction, in which was factored 2) the eyes positions in the field of view (FOV). The points of gaze were represented in a visual panorama (middle panel), with a 10° resolution. The geometric forms on the top represent the position of the landmarks in the panorama, and the brown lines the position of paths. Cells activities were then distributed as a function of visual bins (example with the fainted grey histogram). A 72-degree-period sine-wave (blue) was computed as a reference of a landmark-gazing-responsive cell. Finally, the offset between the neuron’s activity and the reference sine-wave was computed (right panel). B-G: Map of activity as a function of monkey’s gaze position into the visual panorama for 6 PPC single unit examples (the same as in Figure 1F-K). Note that, for a clearer vizualisation of activity modulations, the ordinate axis scale varies across individual cells’ plots, and may not necessarily begin from zero. The correlation coefficients (r) and p-values of the Pearson correlation tests between the averaged activity and the reference sine-wave are indicated for each cell. H-M: Same as B-G, for hippocampal cells, corresponding to the neurons in Figure 1L-Q. N: Correlation coefficient of all PPC (left, N=111) and HPC (right, N=111) cells. The gaze-related activity of 28/111 (25.2%) of PPC cells (red) and of 9/111 (9.1%) of HPC ones (green) were significantly positively, or negatively correlated with the reference sine-wave (colored markers), and overall the average absolute coefficient was higher in PPC than in HPC (µ|r|=0.26±0.035, µ|r|=0.19±0.026; Wilcoxon rank sum: p=0.0072). O-P: Averages of the normalized activity of PPC (N=58, in red) and HPC (N=60, in green) landmark cells, as a function of monkey’s gaze position into the visual panorama. They were significantly positively correlated with the reference sine-wave (PPC: Pearson correlation: r=0.92, p=3.48×10^-10^; HPC: r=0.81, p=1.58×10^-6^). Q-R: Averages of the normalized activity of PPC (N=53, in red) and HPC (N=51, in green) paths cells, as a function of monkey’s gaze position into the visual panorama. They were significantly negatively correlated with the reference sine-wave (PPC: r=-0.85, p=1.43x10^-7^; HPC: r=-0.67, p=3.86×10^-4^). S-T: Distribution of the phase offsets between the average activity of each landmark cells and the reference sin-ewave, for PPC (N=58, in S) and HPC (N=60, in T). The population of cells with significant correlations between their gaze-related activity and the reference sine-wave are represented in color. HPC cells often presented a 60°-phase-offset between gaze and the reference sine-wave. This represents about 12° of FOV if considering that 2 landmarks are located 72° away from each other. The mean absolute offset was modestly different between the 2 regions (PPC: µ|offsets|=39.4± 9.2; HPC: µ|offsets|=46.8±7.1; tailed-Wilcoxon rank sum: p=0.031). U-V: Distribution of the offsets between the average activity of each paths cells and the reference sine-wave, for PPC (N=53, in red) and HPC (N=51, in green). W-X: Averages of the normalized neural place maps of PPC (N=58, W) and HPC (N=60, X) landmark cells (see Figure 1). The two maps were significantly correlated (Pearson correlation: r=0.28, permutation test: p=0.0060). Y-Z: Averages of the normalized neural place maps of PPC (N=53, Y) and HPC (N=51, Z) paths cells. The two maps were not significantly correlated when taking in account the activity in the reward path (Pearson correlation: r=0.13, permutation test: p=0.16). In both regions, the place maps of landmark cells tended to be negatively correlated with the path cells one (PPC: Pearson r=-0.16, permutation test: p=0.086; HPC: r=-0.28, permutation test: p=0.002).

Then, we represented the neural activity as a function of gaze position in the visual panorama. The analyses of this section included the 111 PPC neurons, as well as 111 of the 142 neurons from HPC. Figure 3B-M shows the firing rates of the 12 examples of PPC and HPC cells from Figure 1B-M, as a function of the monkeys’ point of gaze on the visual panorama, binned by 10° of FOV. Neurons responded when landmarks were directly gazed at (Figure 3B,C,H,I) or when they were at the periphery of the visual field, i.e. when maze paths were fixed (Figure 3D,E,J). To quantify this modulation by gaze, we used a 72°-period sine-wave, with peaks aligned with the 4 usually visible landmarks in the panoramic scene (Figure 3A, dark blue sine-wave), and calculated the Pearson correlation between this model and each neuron’s firing rate as a function of gaze position. The obtained coefficients (Figure 3N) varied from positive (Figure 3O-R top) to negative (Figures 3O-R, bottom) values. A positive coefficient indicates that the neuron responded preferentially to the fixation of landmarks, and a negative one that it exhibited selectivity for path fixations (which also correspond to the presence of landmarks in the periphery of the FOV). A high absolute value of the coefficient suggests that the cell was sharply tuned to the animal’s fixation of the exact positions of all landmarks or paths. On the other hand, a low absolute coefficient implies either a broad tuning for all the cues, or a selectivity for a single one. The cells were equally distributed between the “landmark cells” (52.3% parietal cells and 54.1 % of hippocampal cells) and the “path cells”. The continuous distribution between the two extremes, suggested a continuous representation of the FOV, with response fields shifting from foveal (landmark cells) to peri-foveal cells (path cells). At the anatomical level, we found no relationship between the depth of cells along the sulcus and the nature of the cells’ activity suggesting that selectivity to landmark and path were expressed equally in LIP and VIP (Figure S3A; Pearson correlation: r_monkey K_=0.13, p_monkey K_=0.32; r_monkey S_=-0.16, p_monkey S_=0.25), and preventing the clear identification of the neurons of these two regions. There were significantly more PPC cells (28/111, 25.2%) with a significant correlation to the 72°-period wave (permutation test, p≤0.05; represented by filled circles in Figure 3N) compared to HPC cells (9/111, 9.1%; homogeneity Pearson’s Chi-squared with Yates’ correction: X²_1df_ =10.51, p=0.0012), and the distribution of correlation coefficients of PPC cells was higher compared to HPC ones (µ|r|_PPC_=0.26±0.035, µ|r|_HPC_=0.19±0.026; Wilcoxon rank sum: Z=2.69, p=0.0072 on absolute coefficients). This was further confirmed when we examined the magnitude of modulation of landmark gazing on neurons’ activities (Modulation coefficient, M, see Methods), as there was a stronger effect in PPC (µM=0.13±0.013) compared with HPC (µM=0.10±0.016; Wilcoxon rank sum: Z=2.42, p=0.016). Thus, PPC cells responded more strongly to retinal stimulations linked to foveation of the maze’s visual panorama than HPC cells. Moreover, a two-way-ANOVA on the neurons’ firing rates revealed a significant effect of the cell “type” (i.e. landmark or path cells; F_1df_ =6.48, p=0.012), and of the neural area (F_1df_ =9.23, p=0.0027) factors, and that those two factors tended to interact (F_1df_ =3.74, p=0.054). Precisely, landmark cells (µFR=5.75±1.74 sp/s) fired more than path ones (µFR=3.28±0.80 sp/s), parietal cells (µFR=6.14±1.79 sp/s) fired more than HPC ones (µFR=3.043±0.84 sp/s), and the effect tended to be amplified for the parietal landmark neurons (for the detailed listing of all the averaged FR of all categories, see Table 1).

**Table 1.** Comparison of the firing rates of PPC and HPC landmark and path cells. Contingency table indicating the average firing rates of the neurons depending on their « type » and their region. Landmark cells fired more than path ones (two-way-ANOVA: F_1df_ =6.48, p=0.012), parietal cells fired more than HPC ones (F_1df_ =9.23, p=0.0027), and the effect tended to be amplified for the parietal landmark neurons (F_1df_ =3.74, p=0.054).

|  | PPC (N=111) | HPC (N=111) | all areas (N=222) |
| --- | --- | --- | --- |
| landmark cells (N=118) | 8.27 ± 3.24 sp/s | 3.32 ± 1.10 sp/s | 5.75 ± 1.74 sp/s |
| path cells (N=104) | 3.81 ± 0.93 sp/s | 2.71 ± 1.30 sp/s | 3.27 ± 0.80 sp/s |
| all cell types (N=222) | 6.14 ± 1.79 sp/s | 3.043 ± 0.84 sp/s |  |

Next, we aimed to define the relationship between stimulus retinal position and the activity of landmark and path cells. The presence of extra-foveal receptive fields in some of the PPC neurons would results in a left or right shift in their firing rate as a function of the fixation point, relative to the landmark, such as visible in Figures 3G and L. In these examples, the responses were higher when animals fixated on the right side of a landmark, which suggests that cells had a non-foveal left receptive field which would correspond to the contra-lateral side of the recording site. In order to assess to what extent PPC and HPC neurons presented such offset in activity, for each neuron we computed the phase offset between their gaze-related activity waves and the reference sine-wave (see Methods, Figure 3A, right panel). Positive phase offsets correspond to an activity concomitant with fixating at the right of the visual cue, and negative ones to left of the cue (Figures 3S-V). The distributions of the phase offset magnitudes for path cells were not significantly different between PPC and HPC for path cells (µ|offsets|_PPC_=56.50±11.12°, µ|offsets|_HPC_=50.24±9.42°; Wilcoxon rank sum: Z=0.52, p=0.60, Figure 3U-V), but they were for landmark cells (µ|offsets|_PPC_=39.38±9.17°, µ|offsets|_HPC_=46.77±7.09°; one-tailed-Wilcoxon rank sum: Z=-1.86, p=0.031, Figure 3S-T). Indeed, more parietal neurons responded when landmarks were directly fixated (Figure 3S), than hippocampal ones, which preferentially responded when landmarks were off-centered by approximatively 12° from the fovea (see phase-to-offset conversion in Methods; Figure 3T). Interestingly, the population of cells that were significantly modulated by the fixation of landmarks (in red on Figures 3N and S) exhibited a preference for eye position located on the left side of those visual cues (µoffset=-18.57; one-sample Wilcoxon signed-rank: W=9, p=0.016). This could be explained by a preference of cells for contralateral eye positions and saccades. However, due to the highly dynamic nature of the visual stimulations deriving from the task, numerous confounding factors co-exist and make the interpretation of such results challenging.

To examine the relationship between the view responses and the virtual position, we computed the average neural place map for the PPC or HPC populations previously parted in two, as shown by Figure 3W-Z. This classification immediately reveals clear distinct spatial patterns for landmark or path cells. Both parietal and hippocampal landmark cells fired as monkey were located in the inbounds, but parietal path cells preferred the rewarded path, while hippocampal ones were more active in beginning of the outbound paths, following the reward delivery. Accordingly, within regions, place maps for landmark and path cells tended to be negatively correlated, in both PPC and HPC (Pearson r_PPC_=-0.16, permutation test: p=0.086; r_HPC_=-0.28, p=0.002). Across brain regions, place maps of PPC and HPC landmark cells were significantly correlated (r=0.28, p=0.006), while place maps for path cells were not (r=0.13, p=0.15). However, when not taking into account for the computing of the correlations the activity in the rewarded path, that, as mentioned above, is the main spatial feature specifically displayed by the PPC path cells, (see Figure S3B), the place maps of the PPC and HPC path cells were then positively correlated (r=0.20, p=0.038). This suggests that the high parietal firing rate in the rewarded path was likely due to internal motivational reward signals, and that otherwise, parietal and hippocampal path cells shared similar activity patterns. When considered in isolation, the response of the parietal path cells when the animal was located in the rewarded path, facing north, may seem inconsistent with their tuning to the fixation of all the maze paths. However, this result was made possible due to the fact that this pool of neurons was additionally active in the center of the maze, where camera rotations rendered the paths visible within the FOV.

To assess the similarity between the parietal and the hippocampal landmark cells, we examined their activity as a function of the animal’s position along paths for all inbound or all outbound ones averaged together (see Methods, Figure S3C and D top panels). The figure represents the landmark or path cells population activity, with cells ordered as a function of their peaks’ position on the inbound or outbound paths. This revealed that PPC landmark cells were more active at the end of the inbound paths while HPC ones had more distributed peaks along those segments (Kolmogorov-Smirnov test on the distribution of the peak position: D=0.27, p=0.025; Figure S3C). Distribution of peak position in the rewarded path and the outbound paths were also different between the path cells of the two areas (D=0.26, p=0.043; Figure S3D, top row), but this was again due to the difference of activity in the path leading to the reward: when considering only the outbound paths, the two areas presented similar place-related activity patterns (D=0.16, p=0.45; Figure S3D, bottom row). The results thus suggest that reward expectancy strongly modulated parietal’s activity, leading to a higher activity along the specific path leading to reward.

In sum, the distribution of responses depending on animal’s fixation of landmarks or paths accounts for a continuous range of selectivity depending on visuo-spatial layout, but parietal cells were generally more strongly modulated by the content of the view than hippocampal ones. The latter tended to be active when the cues were at the periphery of the fovea rather than at its center. Importantly, in both regions, cells were recruited in different virtual positions affording distinct views of landmarks or paths. Notably, we identified substantial differences between parietal and hippocampal position-related activity patterns, suggesting a different task-based processing across regions despite similar views.

### Temporal neural dynamics relative to landmarks appearance reveal anticipatory activities in PPC and HPC

As a cognitive map of the scene may lead to anticipation of landmarks appearance, we investigated the temporal dynamics of neurons’ activities immediately preceding and following the appearance of the landmarks on screen, i.e. in the animal’s FOV (see Methods). In the parietal cortex, 46.9% of the cells (52/111) responded significantly to the appearance of landmarks in the FOV (see Methods; see examples in Figure 4A-B). In hippocampus, the proportion was similar, with 55/137 cells (40.2%; homogeneity Pearson’s Chi-squared with Yates’ correction: X²_1df_ =0.87, p=0.35; Figure 4C-D). The task unfolded in such a way that most of the time, a landmark appeared on the screen concomitantly to the disappearance of the adjacent one. Thus, to check that the activities of the neurons were not actually a response to these disappearances, we isolated the parts of the maze in which solely one landmark appeared in the FOV, i.e. the beginnings of the outbound paths. Even in these conditions, we still found 48/111 (43.2%) of PPC and 42/137 (30.7%) of HPC cells that significantly responded to landmarks appearances.

**Figure 4.**
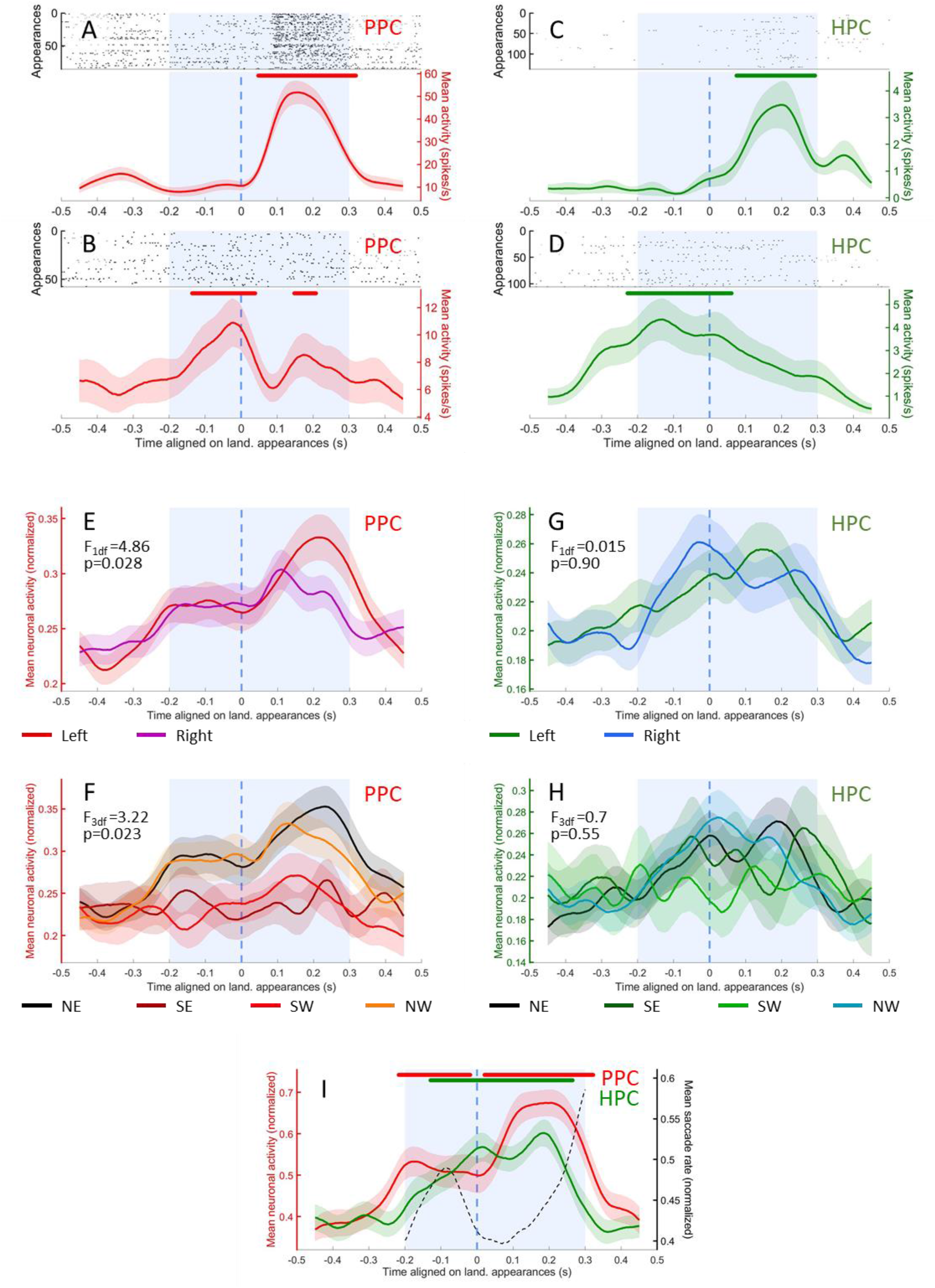
Activity as a function of landmark appearance. A-B: Raster plot (top) and average activity (bottom, in red) of two example PPC cells showing a significant response when aligned on landmarks appearances (blue dashed line), for any positions in the maze. The standard errors are indicated in light color. The analysis window (- 200 to +300ms relative to appearance) is represented in a blue shaded area. The line above indicates times for which activity was significantly higher than baseline (for A: from 47 to 319 ms; for B: from -137 to 39 ms and from 147 to 208 ms). C-D: Same as A-B, for two hippocampus example cells. The activity was significantly higher than baseline from 74 to 293 ms relative to appearance for C, and from -229 to 61 ms for D. E: Averaged normalized activity of the PPC cells responsive to landmark appearance (N=52), aligned on the landmark appearance times (blue dashed line), taking into account every position in the maze. The peri-appearance activity is separated by the side of appearance of the landmarks in the field of view (left in red, right in purple). The standard errors are indicated in light color. The analysis window (-200 to +300ms relative to appearance) is represented in a blue shaded area. The population displayed a significant difference in response rate between the two conditions (3-way-ANOVA: F_1df_=4.86, p=0.028). F: Same PPC averaged activity as in E (N=52), with peri-appearance activity separated by the identity of the appearing landmark (north-east in black, south-east in burgundy, south-west in red, north-west in orange). The population displayed a significant difference in response rate between the four conditions (3-way-ANOVA: F_3df_=3.22, p=0.023). G: Same as in E, for HPC cells responsive to landmark appearance (N=55), with peri-appearance activity separated by the side of appearance of the landmarks in the field of view (left in green, right in blue). There was no significant difference in the population’s response rate when comparing these two conditions (3-way-ANOVA: F_1df_=0.015; p=0.90). H: Same as in F, for HPC cells responsive to landmark appearance (N=55), with peri-appearance activity separated by the identity of the appearing landmark (north-east in black, south-east in dark green, south-west in light green, north-west in blue-green). There was no significant difference in the population’s response rate when comparing these four conditions (3-way-ANOVA: F_3df_=0.70, p=0.55). I: Averaged normalized activity of PPC (N=52, in red, left Y axis) and HPC (N=55, in green, left Y axis) responsive cells aligned on the landmark appearance times (blue dashed line). The dotted black line represents the rate of saccade starts (right Y axis). The standard errors are indicated in light color. The analysis window (-200 to +300ms relative to appearance) is represented in a blue shaded area. The line above indicates time for which activity was significantly higher than baseline (for PPC: from -218 to -19 ms and from 20 to 322 ms; for HPC: from -130 to 265 ms).

Then, we determined the selectivity of individual cells depending on side of appearance of the landmarks (left or right), or their identity (north-east, south-east, south-west or north-east). Importantly, the identity factor here comprises both the visual features of the object, and their position in the maze, since for each session, one landmark will still remain at the same location in the spatial layout. Two-way-ANOVAs (see Methods; p≤0.05) revealed that 26/52 (50.0%) of the parietal responsive cells were selective to the side, and 23/52 (44.2%) to the identity, with 18 (34.6%) of them being selective to both. In hippocampus, only 8/55 (14.6%) of the responsive cells were selective for one side over the other, and 9/55 (16.4%) for the landmark identity, 3 of them (5.5%) being selective to both factors. There were as many cells that preferred the left side compared to the right side in PPC (16/26, 61.5%; homogeneity Pearson’s Chi-squared with Yates’ correction: X²_1df_ =0.31, p=0.58) and in the hippocampus (4/8, 50.0%). However, at the population level, a 3-way-ANOVA on the cells’ averaged normalized firing rate (fixed factors: side of appearance of the landmarks and identity, random factor: cell identity, see Methods) revealed that parietal cells were more active when landmarks appeared on the left of the screen (F_1df_=4.86, p=0.028; Figure 4E). As the recording were all performed in animal’s right hemispheres, the results are consistent with a preference for the contralateral FOV in the parietal cortex. Additionally, the cells were more active when the north-west or the north-east landmarks appeared on screen (F_3df_=3.22, p=0.023; Figure 4F), which correspond to the landmarks that surrounded the reward location. In the hippocampus, there was no preference, neither for the side (F_1df_=0.015; p=0.90, Figure 4G), nor for the landmark ID (F_3df_=0.70, p=0.55; Figure 4H).

Finally, we examined the temporal dynamics of the population activity relative to the landmark appearance across regions (Figure 4I). The averaged activity of the PPC responsive cells was higher than baseline (see Methods), from -218 to -19 ms and from 20 to 322 ms relative to landmarks appearance, although the activity peak was post-appearance. Similarly, the hippocampal responsive cells increased their firing from -130 ms pre-appearance, and decreased around 265 ms post-appearance. Hence, the temporal dynamics of both the parietal and the hippocampal cells show an anticipatory activity of the entrance of the visual cues in the field of view, with the parietal cortex slightly preceding hippocampus. It is noteworthy that the rise in PPC activity also anticipates the saccade rates (black dashed line), suggesting a memory guidance of directed saccades.

Overall, our results show that parietal and hippocampal neurons are sensitive to the appearance of meaningful visual cues in the field of view, and encode spatial features, such as their side of appearance or their position (or identity) in the environment. At the population level, anticipation of landmarks appearance, together with a higher activity for landmarks surrounding the reward location in parietal cortex, suggest a representation of the spatial layout in a cognitive map, with objects’ saliency established according to the internal agenda.

## Discussion

To understand the neural basis of vision-based navigation, we compared neural activity in intraparietal areas to that of the hippocampus, while two monkeys searched a virtual star-maze for a hidden reward. More parietal neurons were modulated by animal position in the virtual maze than hippocampal ones, and they expressed more stereotyped patterns than in the hippocampus (Figure 1). Active visual exploration during navigation was accompanied by saccade-related neural activity; this was expected in the parietal cortex, but was also observed to a lesser extent in the hippocampus. Nevertheless, oculomotor activity *per se* did not solely account for spatial selectivity in either area (Figure 2). Parietal activity especially, was strongly linked to the fixation of visual elements of the environment such as landmarks or paths, thereby resulting in specific virtual-position selectivity (Figure 3), connecting view and task-specific contexts. The analysis of the temporal neural dynamics of cells relatively to landmarks appearances in the field of view suggested that both regions anticipated them, but only parietal neurons were modulated by landmark’s proximity to the reward (Figure 4). These results show how parietal cortex and hippocampus support a continuous representation of visual space, through dynamic and successive recruitment of neurons, during action preparation and acquisition of elements in visual space. The results shed light on how seemingly place codes bind views and actions in space with an internal task-based map.

### Interpreting position selectivity

Virtual navigation resulted in a greater recruitment of parietal neurons than hippocampal ones, and further, their selectivity was as strong as in hippocampus – yet qualitatively distinct. Thus, virtual-reality-generated visual stimulation resulted in parietal spatial selectivity that could be interpreted as “place” codes akin to those found in rodent hippocampal place cells. The only significant difference consisted of parietal neurons displaying repeated patterns of position selectivity, for similar subsegments of the maze. This echoes results showing that rodent posterior parietal cells demonstrates strong pattern recurrence, for similar action-types or postures, at different locations of a real or virtual environment (Harvey et al. 2012; Lee et al. 2022; McNaughton et al. 1994; Nitz 2006; Shelley et al. 2022; Shelley and Nitz 2021).This suggests that parietal cells are recruited for specific types of movements, actions or sensori-motor patterns, linked to specific navigational context. The interpretation of our data may follow a similar line. For instance, the PPC neurons’ enhanced activity before the maze center or at the beginning of the inbound paths (see Figures 1B-G and 3W and Y) may be related to anticipatory or concurrent responses to the onset of rotational or linear movements, as previously observed in rodent PPC (Wilber et al. 2014). Further, in the primates, including humans, parietal cortex supports visuo-spatial attention and eye-movements directed in space (Bendiksby and Platt 2006; Bremmer 2005; Colby and Duhamel 1991; Duhamel et al. 1992, 1998; Goldberg et al. 2006; Kravitz et al. 2011; Platt and Glimcher 1999; Steenrod et al. 2013; Zhang et al. 2014; Zhou et al. 2018), which are functional properties likely recruited during virtual navigation. This could account for the very strong parietal activity observed before the decision point, or in the rewarded path, before the reward delivery, seemingly tiling the space in a function-driven manner. We suggest that the apparent parietal position code can be linked to visual processing of relevant cues, contributing to a task-relevant representation of visual space, rather than pure *place* code.

Indeed, in both regions, the nature of cues processed during fixations impacted the cells strongly. Neurons displayed a range of responses to landmarks and paths at different retinal positions, associated to different virtual positions (Figure 3 and S3). Given the environment layout, the nature of the selectivity of the path cells could arise from a peripheral retinal stimulation by the landmarks or from a central stimulation by the path, the latter being consistent with a role of parietal cortex in encoding route affordance (Arbib 1997; Calton and Taube 2009). Cells responding to landmarks views displayed a higher activity than path cells. This suggests that the direct foveation of cues that are relevant to navigation due to their stable and orientation-informative nature elicits more activities than routes, that geometrically structure the environment in an ambiguous (because symmetrical) fashion. Compared to hippocampal cells, parietal activity was more strongly controlled by visual cues, as expressed by 1) a stronger correlation between rate as a function of point of gaze and the visual scene layout (Figure 3N), 2) a higher modulation coefficient, and 3) a majority of responses to landmark-centered gaze (Figure 3S). By contrast, in the hippocampal cells sensitive to landmarks viewing, the relationship between the neuron’s rate as a function of gaze and the visual scene layout was generally lower, and maximized when an offset was introduced between them (Figure 3T). This result echoes previous findings, reported in freely moving animals, in which hippocampal cells encoded facing location with no to little added value of direct gaze (Mao et al. 2021), suggesting that what is in the view is more important than the precise gaze position. Our results show that the previously identified link between hippocampal activity and view (Corrigan et al. 2023; Doucet et al. 2020; Hoffman 2013; Jutras et al. 2013; Wirth et al. 2017) is weaker and more indirect than in the parietal cortex, the latter being more driven by features of visual input. Thus, although hippocampus receives input from the parietal cortex (Kravitz et al. 2011; Rockland and Van Hoesen 1999; Selemon and Goldman-Rakic 1988), its activity does not simply reflect the presence of a spatially or behaviorally informative object in the visual receptive field, but integrates other factors of the task.

An interesting picture arose in the two regions, when we computed the average neural place map for landmark or path cells, revealing how position-related firing patterns are linked to visual cues and their task-relevant processing (Figure 3W-Z). In line with a role of parietal cortex in encoding animal’s motivational state (Bendiksby and Platt 2006; Leathers and Olson 2012; Platt and Glimcher 1999), parietal “path” cells displayed a higher activity in the path leading to reward. By contrast, hippocampal cells were more active following the reward delivery, in line with previous reports showing that hippocampus encodes reward outcome in primates (Brincat and Miller 2015; Camara et al. 2009; Corrigan et al. 2022; Knudsen and Wallis 2021; Wirth et al. 2009; Wittmann et al. 2005) and reward locations in rodents (Dupret et al. 2010; Hok et al. 2007; Hollup et al. 2001; Markus et al. 1995). For landmark cells, the highest modulation occurred in parietal cortex when animals were close to decision points in inbound paths, while hippocampal activity occurred earlier in those segments (Figure 3W-X), suggesting that each region takes part in different processes. Specifically, the spatial selectivity of parietal cells suggests it may take part in action-guidance and execution such as saccades or joystick moves controlling animal’s virtual motion, while hippocampal cells may play a role with landmark identification. Despite their superficially similar responses to actions such saccades or cues fixations, the general sensorimotor context results eventually in a distinct, complementary task-based representation of the space between the two areas.

### Contextually-modulated saccade-related activity

Saccades, which are characteristic of visual exploration, bring elements of the visual scene into focus. Consequently, they can serve as a proxy for visual attention (Gottlieb et al. 2014, 2014, 1998; Hayhoe and Ballard 2005; Meister and Buffalo 2016; Rucci and Poletti 2015; Zhu et al. 2022) and play a part in the subsequent formation or consolidation of memories (Henderson et al. 2005; Loftus 1972; Melcher 2001; Olsen et al. 2016; Pertzov et al. 2009). Accordingly, monkeys made many saccades when landmarks appeared in the visual field, or before decision points (Figure S2), and these were followed by fixations of landmarks and paths. This active exploration likely reflects information-seeking that guides navigation (Lakshminarasimhan et al. 2020; Zhu et al. 2022). Concomitantly to this explorative behavior, we observed a strong recruitment of the intraparietal neurons during saccades. The observed pre-saccadic increase was characteristic of saccade-preparation and target-selection activities, described in LIP cells (Bendiksby and Platt 2006; Goldberg et al. 2006; Grefkes and Fink 2005; Steenrod et al. 2013). Further, following a previously described trans-saccadic suppression (Bremmer et al. 2009; Bridgeman et al. 1975; Herrington et al. 2009; Ibbotson et al. 2008), many cells displayed the post-saccadic activity enhancement, generally observed in VIP or LIP when a stimulus enters a foveal or peri-foveal receptive field, concomitantly to saccade end (Ben Hamed et al. 2001; Duhamel et al. 1998; Munuera and Duhamel 2020; Zhou et al. 2018). Consistent with previous work underscoring the difficulty of isolating the saccade response from the visual stimulation response (Bremmer et al. 2009), we could not clearly separate LIP from the VIP along the depth of the sulcus using cell’s activity profiles (Figure S2N and S3A). This indicates that the rich visual stimulation of the task recruited cells in both regions, with properties either functionally overlapping or expressed for co-occurring features. Previous works suggests that the post-saccadic enhancement described here, would be a global, active phenomenon, spreading from V1 across the whole visual cortex, whose neural activities synchronise with saccades rhythm to optimize the visual treatment of the foveated stimuli (Ibbotson et al. 2008; Ito et al. 2011; Rajkai et al. 2008; Zanos et al. 2015). In line with this, the task context, embodied in virtual position, influenced both baseline activity and the amplitude of the saccade response itself in the parietal cortex (Figure 2). Moreover, at the population level, there was no preference for one part of the maze compared with others, suggesting that decisional, attentional or motivational context weighted on the strength of the baseline signal in which saccades to landmarks were made in a complex way, differing across the cells.

Surprisingly, we found that a small population of hippocampal cells also displayed perisaccadic activity, with a similar distribution to that of parietal perisaccadic responses (Figure 2H), including pre-, trans-, and mostly post-saccadic preferences. These results are consistent with the previous findings linking hippocampal activity not only to view but also to eye movements (Doucet et al. 2020; Liu et al. 2017; Mao et al. 2021; Ringo et al. 1994; Ryals et al. 2015; Sobotka et al. 1997) and confirm the importance of active visual exploration for hippocampal processing (Corrigan et al. 2023; Doucet et al. 2020; Hoffman 2013; Zhu et al. 2023). The results document for the first time how a naturalistic task strongly engages neurons within the intraparietal areas and hippocampus, and how their activities reflect the impact of task-context on oculo-motor control, during goal-directed navigation.

### Processing and anticipating landmarks across regions

Our results showed that, in accordance with what is showed in literature, parietal neurons responded to the entrance of a moving object in the field of view (Bremmer et al. 1997, 2002b; Duhamel et al. 1998). Further, that selectivity to landmark was present in individual cells beyond the simple contralateral preference often described (Duhamel et al. 1998; Sereno et al. 2001; Wardak et al. 2002), although the whole population did show a preference for the contralateral hemifield. While such selectivity is expected in the hippocampus, parietal cells preferred landmark closer to reward, in line with the notion that posterior parietal cortex processes the motivational relevance of cues and contributes to representing the visual environment through a saliency map (Falkner et al. 2013; Goldberg et al. 2006; Leathers and Olson 2012; Zhang et al. 2014). As we previously showed (Wirth et al. 2017), this preference was not carried over to hippocampal cells, which appeared to represent landmarks more evenly as a population, despite selectivity being observed at the individual level. This observation is congruent with an encoding of landmarks in the HPC as topological cues that structure the environmental layout, and separately provide orientation-relative information. We may hypothesize that, while PPC would rather encode the *position* of the relevant visual cues neighbouring reward, the hippocampus would more likely be selective to the *visual identity* of the objects, defined by their qualitative features. Our results are consistent with this hypothesis that parietal cells encode the saliency or relevance of landmarks with respect to the current task goal, while the hippocampus maintains a global internal map of space.

Interestingly, we also found that both parietal and hippocampal neurons anticipated the appearance of a landmark in the FOV. While we previously identified this anticipation in the hippocampus (Wirth et al. 2017), the current results illustrate how parietal activity displays a similar pattern (Figure 4), likely consistent with exploratory saccades targeting upcoming landmarks, and with the role of PPC in such target selection (Bremmer et al. 2016; Hagan et al. 2012; Heiser and Colby 2006; Platt and Glimcher 1999; Steenrod et al. 2013). It has been previously proposed that hippocampus receives an indirect efference copy of the eye movement from the superior colliculus (Katz et al. 2022; Martinez-Trujillo 2022; Ringo et al. 1994; Sobotka et al. 1997). The temporal dynamics of the peri-saccadic response are compatible with this hypothesis, with the presence of some hippocampal pre-saccadic neurons (Figure 2). The hippocampal temporal dynamics are also in line with the notion that the hippocampus serves as a predictive map (Stachenfeld et al. 2017; Zhu et al. 2023), likely anticipating the upcoming landmark. The nature of the parietal temporal dynamics relative to landmark appearance supports the existence of such a predictive coding in parietal cortex as well, allowing controlling exploratory behavior towards anticipated targets continuously linking timely action and objects in space. The phenomenon of *predictive remapping*, through which LIP neurons begin to respond to stimuli presented in their future receptive field prior to saccade execution (Duhamel et al. 1992; Heiser and Colby 2006; Kusunoki and Goldberg 2003; Zhou et al. 2018), implies that the neurons have access at any time to visual information covering the whole FOV, giving an impression of “anticipation”. In addition, the tonic activity of LIP neurons that maintains the location of a recently extinguished stimulus in working memory (Barash et al. 1991; Ben Hamed et al. 2001; Duhamel et al. 1992; Hagan et al. 2012) suggests that they can also represent non-visually-present cues within the FOV. Our results demonstrate that, further, PPC neurons may hold a representation of the environment that extend beyond the FOV. This may be achieved through top-down connection from HPC to PPC (Kobayashi and Amaral 2003, 2007; Rockland and Van Hoesen 1999; Selemon and Goldman-Rakic 1988), that would provide the latter with information extracted from the cognitive map held in memory.

Altogether, the complementary spatial activity maps of the two regions suggests that, despite the superficial similarity of their responses to actions such as saccades or to sensory events such as landmark appearances, their selectivity to specific contexts results in a task-based discrimination of animal’s position in its environment and its orienting towards the goal. Ultimately, those spatial response allows for the linking of action and objects in space and memory. Future work should further explore the impact of the environment familiarity on such spatial encoding, and its evolution through learning stages.

## Supplementary material

## Methods

### Ethics statement

Our study involved two nonhuman primates. All experimental procedures were approved by the animal care committee (Department of Veterinary Services, Health & Protection of Animals, permit no. 69 029 0401) and the Biology Department of the University Claude Bernard Lyon 1, in conformity with the European Community standards for the care and use of laboratory animals (European Community Council Directive No. 86–609). Further, our procedures were examined by CELYNE, the local ethics board, which approved the in vivo methods used in the laboratory. We minimized animal suffering and maintained their well-being by using anesthetics and pain management during surgeries for recording chamber implantation. During the experiments, animal’s behavior and well-being was monitored.

### Behavioral methods and setup (Figure 1)

Animals were head restrained and placed in front of a large screen (152 x114 cm), at a distance of 101 cm, with a FOV of 74cm. They were further equipped with active shutter glasses (Nuvision), coupled to the computer for 3-D projection (DepthQ projector, Infocus) of a virtual world (Monkey3D, Holodia). The projection parameters were calibrated to render objects’ size real by calibrating disparity using the actual interpupillary distance of the monkeys (3.1 cm for monkey K and 3.0 cm for monkey S). We confirmed the animals perceived images with the depth of stereoscopic projection by measuring vergence, as a small object moved from an apparent 50 cm in front of the screen, to 150 cm behind the screen. To this end, two small infrared cameras were mounted above each eye, and the movement of the pupils of each eye was monitored (ASL). The cameras further allowed monitoring the animal’s gaze through the task. Animals learned to navigate via the joystick towards a reward hidden at the end of one of the star-maze paths (Figure 1E, ). The star-maze had a radius of 16m, and speed of displacement was 5m per second. This velocity was chosen to optimize the number of rewarded trials in a session, and prevent the animals from getting too impatient.

During a shaping period that lasted 6 months, animals learned to find the reward targets whilst operating a joystick that controlled a sphere on the screen. Once they had mastered this task, they were introduced to a 3-D-version of this task. Then, they were trained in a simple Y-maze, in which they had to move the joystick to approach the sphere. Next, landmarks were introduced along the Y-maze, and animals were trained with the sphere in presence of the landmarks. Then, the sphere was removed and animals were rewarded when they went toward the end of the arm where the sphere was last. To this end, they had to use the landmarks. At this point, they were finally introduced to the full star-maze. For one animal that would not go to the end of an arm if a sphere was not there, a different strategy was adopted: we replicated the sphere five times and changed the rules such that there was a sphere at each end, but the animal had to find “the one” which would give a reward and blink when approached. Once this step was learned, the spheres were removed for him as well. Finally, animals were trained to learn new landmark arrangements every day.

We used a star-shaped environment rather than using an open field to ensure multiple passes through the same trajectories, and to avoid locations with too sparse data. Each day, the animals had to locate a new position of the reward with respect to new landmarks. Each trial began with the animal facing the maze from one inbound path end. The joystick allowed the animal to move to the center and turn left or right to choose and enter one path. Once the animal reached the end of the path, it was given a liquid reward only if correct, and then brought to a randomly chosen new start, whether the trial was correct or incorrect. Figure 1A presents the sequence of a trial from above (top panel) and from the animal’s perspective (bottom panel).

### Mapping the animal’s point of gaze in the allocentric reference frame (Figure 3)

We computed the point of gaze in an allocentric frame, wherein objects (landmarks or paths) or positions in space towards which the monkey gazed were mapped (Figure 3A). To calculate the point of gaze, we projected the point of gaze on a cylindric wall intersecting landmarks and path (Figure X), using orientation of the camera as a function of its position, and combined with the X and Z eye position in the field of view. The points of regard were mapped onto a vertical circular wall enclosing the landmarks; this wall was then cut and flattened into a visual panorama collapsing the vertical dimension. This panorama represents where the animal is gazing in the spatial scene, not the craniocentric eye position. The coordinates obtained were then used to compute the firing rate of the cells as a function of the animal’s point of regard (Figure 3B-M; see *Correlation between gaze activity and reference sine-wave*).

### Electrophysiological recordings

For a period of approximately 6 months, each animal underwent daily recording sessions, during which laminar U-probes (Plexon®) were lowered to the target areas (see Figure S1A-B for PPC, and (Wirth et al. 2017) for the HPC), both located in the animal’s right hemispheres. Recordings began if individual cells were present on the contact electrodes, and the task was then started. Individual cells were pre-sorted online and re-sorted offline (offline sorter, Plexon Inc.), and only cells whose waveforms possessed reliable signal-to-noise ratios (two-thirds of noise), high enough activity rate (activity peak superior to 1 spikes/s), and whose activity was stable in time for at least 10 trials per starting inbound paths (40 trials in total) were included in the database. One hundred and eleven cells recorded in the intraparietal sulcus were kept for place- and saccades-related analyses, and 142 cells recorded in CA3, CA1 or the dental gyrus of hippocampus. However, some of the eye-tracking data were missing for a population of HPC cells, and therefore, only 111 over 142 HPC cells were used for the gaze-related activity analyses.

### Data analysis

All data were analyzed with custom Matlab scripts. The normality condition of the samples being not always respected, we preferred to use non-parametric statistical tests for most of the analyses, allowing a greater robustness. For all tests, the α risk was set to 5%. Analysis were performed only on the data of the correct trials.

### Wilcoxon rank sum test

As the normality was not respected for most of our sample, and for more robustness, we used Wilcoxon rank sum test when two medians had to be compared. The exact method was used when N_sample 1_<10 and N_sample 1_+N_sample 2_<20, and a normal approximation for large samples.

### Neural place maps & saccades maps

We computed each cell’s mean firing rate for each spatial bin, by simply dividing the number of spikes recorded in that bin by the total time spent in it. Only bins comprising at least four successful trials were kept. For display (Figure 1B-M and S1C-J), a smoothing procedure was applied: the instantaneous firing activity was slightly smoothed with a Gaussian kernel (std=100ms) before computing the map. When comparing spaces, no smoothing was used and bin sizes were adjusted so that each map contained a similar number of bins (Nbin=113). The same protocol was used to produce saccades maps, except that instead of the neuron’s spikes, the animal’s saccades were counted, using the time of their ends. A saccade was defined following Behrens’ adaptive threshold for angular eye acceleration method (2010), with the initial threshold set as 3.4std. Starts and ends of saccades were thereby detected. Because a slight variability in saccade density was noticed across sessions, the correlations tests have been performed between the neural place maps and the saccades maps, both computed for each individual cell. As several cells were recorded on a single session, different cells could share the same saccades map. To study the similarity between two maps, either place or saccades ones, a permutation test was performed: surrogated data were created by randomly shifting one of the two 113-bins-long-matrix 999 times, and Pearson correlation tests were performed for each of the surrogated data, as well as for the actual one. The rank of the actual correlation coefficient among the set of 1000 (actual+999 surrogate ones) was used to extract a statistical p-value (bilateral test). To generate mean maps of a population of cells, the matrices were normalized using standardization method, before being averaged.

### Information Content (IC)

For each individual cell, we iteratively adjusted the spatial resolution of their place map to get as close to 200 valid bins as possible. Bins were considered valid if they included more than 400ms of time in successful trials. We computed the information content in bits per spike with the following formula (Skaggs et al. 1992):

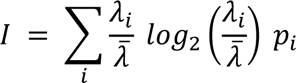

where *λ_i_* is the firing rate in the spatial bin *i*, *λ̅* is the mean firing rate, and *p_i_* is the fraction of the time spent by the animal in bin *i*. IC is zero for a homogeneous firing over the *M* bins; it is equal to *log_2_(M)* when a single bin contains all the spikes and the animal spends an equal amount of time visiting each bin. To avoid potential bias, we normalized the IC by subtracting from it the mean IC of the 999 surrogate datasets: we first created 999 surrogate data sets, in which we divided the recording time into chunks of 5 seconds, that we randomly shifted. This procedure decorrelated the spikes from the animal’s behavior while essentially preserving the structure of spike trains (e.g. spike bursts). The analyses were run on actual and surrogate data, and for any tested variable, the rank of its actual value among the set of 1,000 (actual+999 surrogate ones) was used to extract a statistical p-value (bilateral test).

### Sparsity index (S)

Following standard procedures (Hurley and Rickard 2009; Ravassard et al. 2013), we estimated sparsity by the ratio of L1 norm over L2 norm and defined as sparsity index:

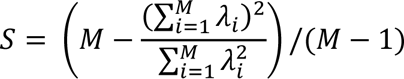

where M is the number of spatial bins and λ*_i_* the firing rate in bin *i* as above. The sparsity index S is 0 for a homogeneous firing map and 1 when a single bin contains all the spikes.

### Depth of tuning Index (DT)

The depth of tuning index was computed as a classic selectivity index:

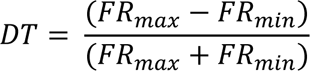

### Characterization of place fields

Place bins with neuronal activity above the threshold of mean + 2std of the activity over the whole maze were selected. Then, they were sorted from higher to lower FR, and the first one was taken as the center of a place field. All the other selected bins within a distance of 5.6 units were considered as part of the same place field than the central one, and were removed from the list. The process was repeated until no bin above threshold was left.

### Symmetry of place-related activity

For each cell significantly modulated by position (see *Information Content*), we compared their mean FR for the right part of the maze with the one if the left part with a two-tailed Wilcoxon rank sum test, considering activity only in inbound paths, outbound paths or both. For the population, we performed a two-tailed Wilcoxon matched pairs signed-rank test, comparing the left *versus* right activity of all cells, in inbound paths, outbound paths or both.

### Kruskal-Wallis for saccades frequencies and durations

For each maze segment (inbound paths, center, rewarded path, outbound paths) of each trial, the number of saccades detected was normalized by the time spent in the segment, to obtain a saccades rate in saccades/seconds per segment, for each trial. The median saccades rates of the 4 segments were compared using a Kruskal-Wallis test, followed by a multiple comparison test, using critical values from Student’s t-distribution, adjusted for multiple comparisons with the Bonferroni method. The same method was applied to compare saccades durations. Saccades durations were computed by subtracting the time of start from the time of end, for each saccade of each maze segment.

### Neuronal activity aligned on saccades ends (perisaccadic activity)

The ends of saccades were used instead of starts because literature showed that they were more informative in the brain, with neurons more precisely tuned to them, as well as sensations and perceptions (Zhou et al. 2018). The spikes of each cells were aligned on each saccade end and distributed in 1ms-long time-bins, so we obtained a 1900ms-by-N_saccades_-long matrix, with the first time-bin corresponding to -899ms relative to saccades ends, and the last one to +1000ms. Then, the spike counts for all the saccades were averaged to obtain a raw mean saccade-relative activity, and finally smoothed using a Gaussian kernel (mean=0, std=35ms). Using a conservative criterion, a cell was considered as saccade-responsive when, considering the saccades from the whole maze, the maximum or the minimum of the smoothed activity in the response window (-150 to +250ms relative to saccades starts, i.e. -242 to +157ms relative to saccades ends) was superior or equal, or inferior or equal, respectively, to the mean ± 3.5std of the baseline (-899 to -242ms and +157 to +1000ms). We specifically chose a higher threshold because activity in response to saccade was more stereotyped, and thus displayed lower intra-cell variation. After that, to create the population activity, each cells’ raw mean activity was at first convoluted with a Gaussian kernel (mean=0, std=50), then normalized using the Min-max normalization, before being altogether averaged. The population activity was considered as significantly responsive to saccades for each time-bin of the response window (-300 to +300ms relative to saccades ends) in which this average value was superior to or inferior to the mean ± 2.5std, respectively, of the baseline (-999 to -300ms and +300 to +1000ms).

### ANOVA on the saccadic responses rate and on their amplitude

To compare the response rate of the neurons to saccades performed from different portions of the maze, for each cell, its raw activity over the whole response window (-150 to +250ms relative to saccades starts) was averaged for each saccade of the maze portion. This resulted in 4 matrices (1 for inbound paths, 1 for center, 1 for rewarded path, 1 for outbound paths) containing 1 value of global activity per saccade. Within each cell, the mean of those 4 matrices were compared using a one-way-ANOVA, and critical values from Student’s t-distribution, adjusted for multiple comparisons with the Bonferroni method for post-hoc analysis. To compare the amplitude of the neurons’ responses to saccades, for each cell, and for each maze segment, its minimal and maximal activity within the response window was identified from its smoothed activity matrix. The mean minimal response for each saccade was the averaged raw activity of a window of -25 to +25ms around the minimal response, and the same protocol was used for the mean maximal response. Finally, for each saccade, the amplitude of the response was determined by subtracting the mean minimal response to the mean maximal response. This resulted in 4 matrices containing 1 value of amplitude per saccade. The mean of those 4 matrices were compared using a one-way-ANOVA with a Bonferroni-corrected Student’s t-distribution-based post-hoc. To determine whether the segment in which a cell exhibited its highest activity coincided with the segment where the animal executed the longest saccades, we assessed, at the population level, whether the observed probality of both the preferred segment and the longest saccade duration occurring in the same segment exceeded chance (chance level 4.5).

### Neurons depth in IPS

The coordinate of each cell was computed by combining the placement of the electrode on the cranial implant with the location of the recording contact on the probe. The inclination of the intraparietal sulcus was determined using the anatomical MRI of the animals, for each frontal slice. Then, each cell’s depth relative to the anatomical dorso-ventral axis was converted to coordinate relative to the sulcus inclination. The zero of the depth axis thereby created correspond to the bottom extremity of the sulcus.

### Correlation between gaze activity and reference sine-wave

The FOV was split in 36 bins of 10° each, centered on the goal, and the point of gaze of the animal was computed (see *Mapping the animal’s point of gaze in the allocentric reference frame*) for each spike of the neuron. After a temporal normalization, those spikes were distributed along the visual bins to obtain a firing rate as a function of animal’s gaze position in the visual panorama. These data were smoothed using a normal probability density function, with a mean of 0, and a standard deviation of 1.2. As only the correct trials were used for the analyses, the monkey never faced the southern landmark. Thus, we removed the corresponding 8 bins out of the 36-bin-long matrix of FR, to be left with a 28-bin-long matrix. Each of the 4 landmarks was separated from the others by 7.2 bins (i.e. 72° of the FOV), and that was also the case for the 5 paths. A reference wave was created, as a sinus function whose period was 7.2 bins, and whose peaks were centered on the landmarks positions. To assess whether the neurons significantly responded to the gazing of landmarks or of paths, a permutation test was performed: surrogated data were created by randomly shifting the cut, smoothed gaze-related activity matrix 1999 times, and Pearson correlation tests were performed between each of the surrogated data and the reference sine-wave, as well as for the actual one. The rank of the actual correlation coefficient among the set of 2000 (actual+1999 surrogate ones) was used to extract a statistical p-value (bilateral test). If the correlation coefficient (r) was positive, the cell was considered as a “landmark cell”, and if it was negative, the cell was considered as a “path cell”.

### Modulation coefficient (M) and offsets with reference wave

The modulation coefficient was here computed as the magnitude of the signal of the smoothed gaze-related activity at the frequency of the reference sine-wave, divided by the sum of the magnitudes at all frequencies found in the smoothed gaze-related activity, determined by Fast Fourier transform (M.Sc. Eng. Hristo Zhivomirov, 2014). In other words, it assesses how strongly the gaze-relative activity is oscillating at the frequency of our reference sine-wave, relatively to all of the other frequencies present in the signal, i.e how much the neurons’ activities were driven by the gazing of the landmarks. Subsequently, the offset was computed, as the shift in space existing between the smoothed gaze-related activity at its main frequency, and the reference wave.

### ANOVAs comparing the gaze populations’ FR

For each cell, we computed their average FR from their smoothed gaze-related activity. Then we tested the effect of area of recording (PPC or HPC) and cell “type” (landmark or path cell) on the FR by performing a two-way-ANOVA, with interaction factor. If a p-value was significant, multiple comparison tests were subsequently performed, using Tukey’s honestly significant difference criterion.

### Comparison of place activity in the different maze segments

The neural place maps matrices of PPC and HPC were cut in order to keep only segments of interest: for the landmark cells, the last 4 bins of the outbound paths and the whole inbound paths; for the path cells, the rewarded path plus the first 12 bins of the outbound paths, and the 12 first bins of outbound paths alone. Then the cut matrices were normalized using the Max-Min method, and averaged, to obtain a mean, linearized population activity.

### Kolmogorov-Smirnov tests on population maps

For the saccade-related, the place-related and the foveation-related activities, comparison of the PPC and HPC cells distributions were performed. For the figure, maximal value of the activities of the selected cells were found, and the cells sorted accordingly, from earlier to later peak. The distribution of those peaks were compared between PPC and HPC using a two-sample Kolmogorov-Smirnov test.

### Neuronal activity aligned on landmarks appearances

The same protocol was used than to align spikes on saccades ends, except that appearance times were used. As the relationship between the camera orientation and position in the maze was consistent across trials and sessions, we first determined which positions corresponded to the entrance of a landmark in the FOV, on average (the different objects could slightly vary in size). Then, the appearance times were defined as the timestamps at which the camera was at those positions. A cell was considered as responsive to landmarks foveations when the smoothed activity in the response window (-200 to +300ms relative to appearance) was superior or equal to the mean + 2.5std of the baseline (-799 to -200ms and +300 to +800ms) for at least 30 consecutive ms. The normalized population mean activity was considered as significantly responsive to landmarks appearance for each time-bin of the response window (-200 to +300ms relative to appearance) whose value was superior to the mean + 2.5std of the baseline (-999 to -200ms and +300 to +1000ms).

### ANOVA on the response rate to landmark appearance

To compare the response rate of the neurons to landmark appearance, we proceeded the same way as when we compared the saccades-response rate (see *ANOVA on the saccadic responses rate and on their amplitude*), at the population level. The average rates of the raw activity in the whole response window (-200 to +300ms relative to saccades appearance) of every responsive cell were compared by the use of a 3-way-ANOVA, including the side of appearance factor (i.e. left or right), the landmark identity (i.e. north-east, south-east, south-west or north-west), as well as a within cell factor.

## Figures

**Figure S1.**
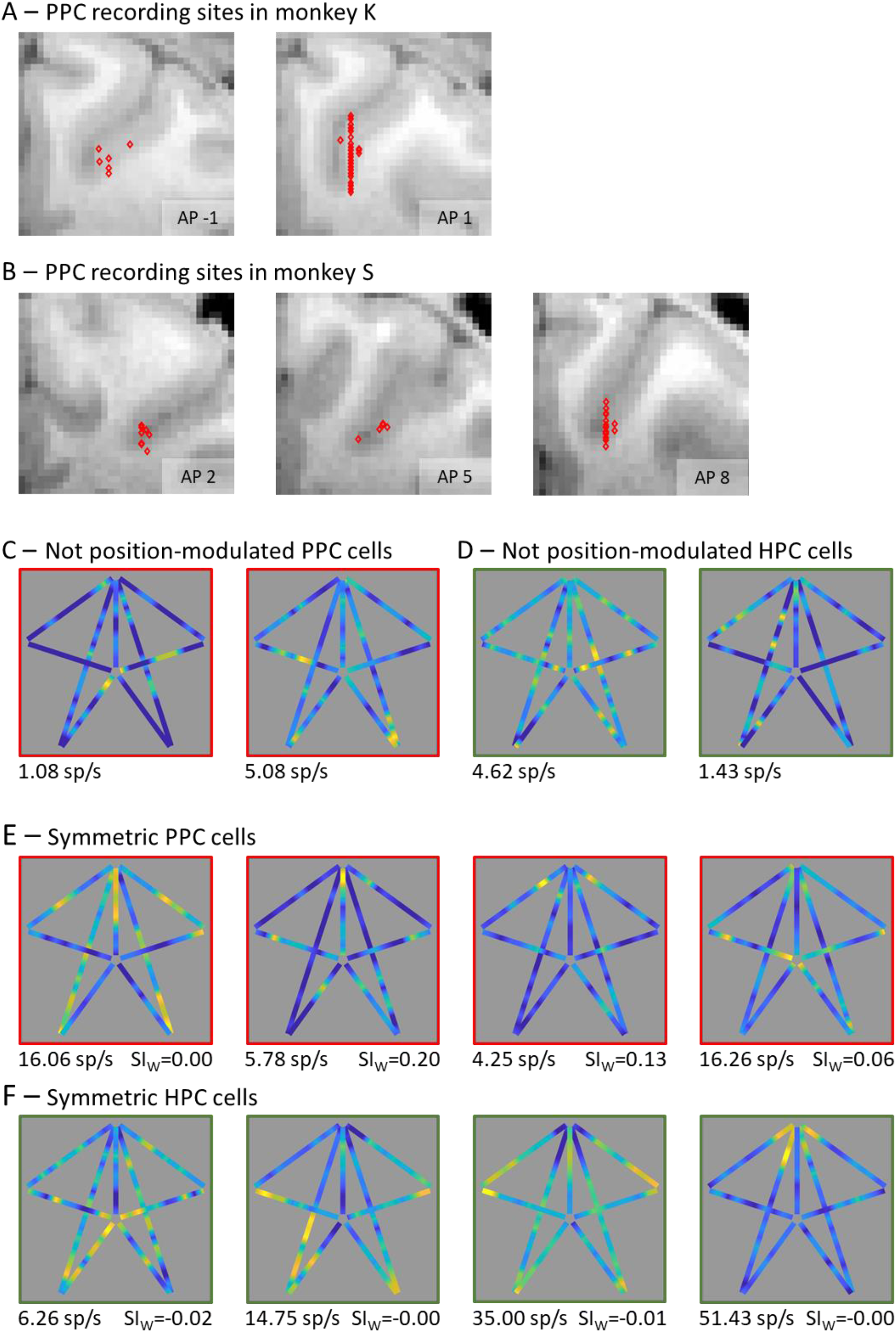
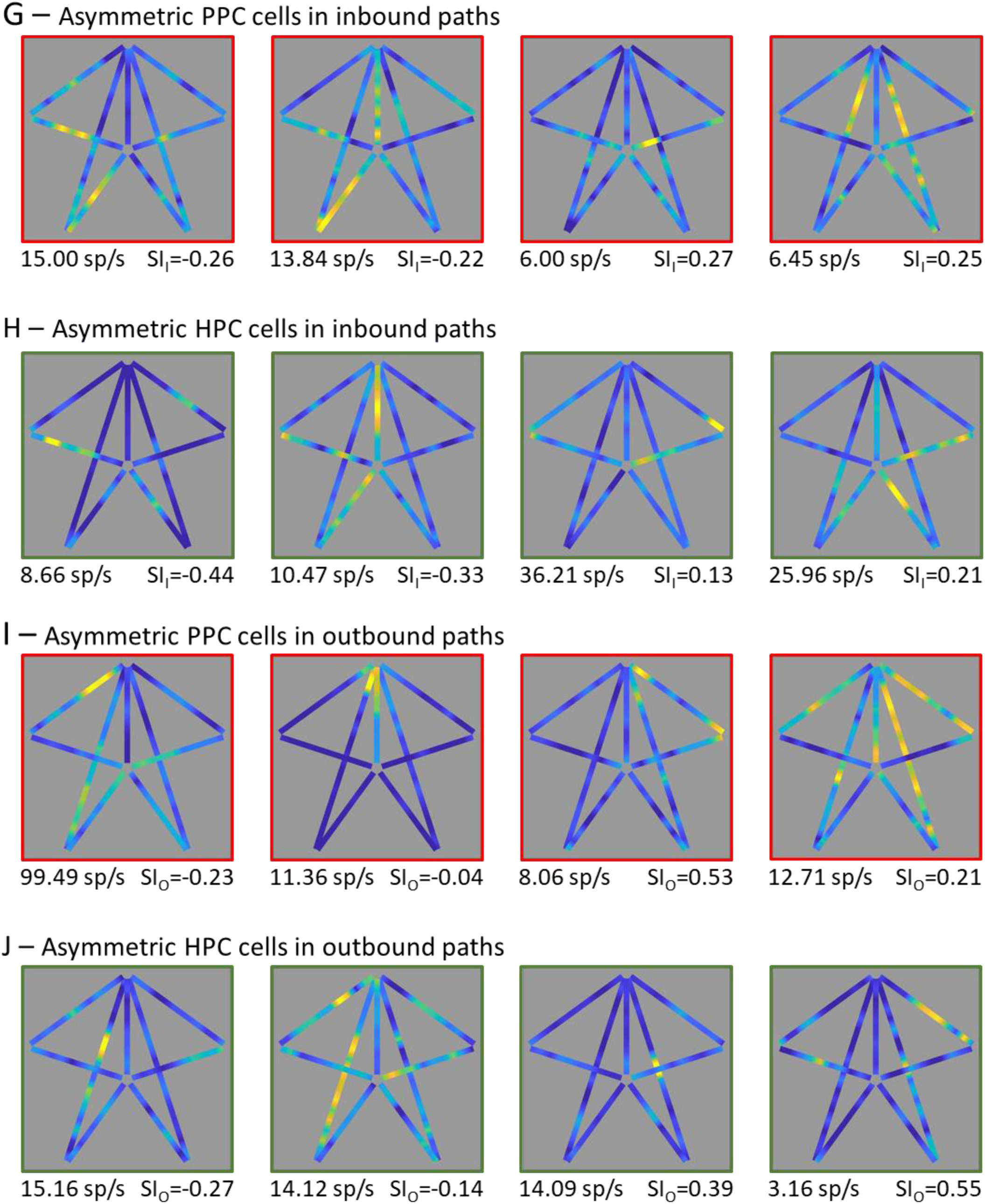
Detailed recording sites and spatial properties of parietal and hippocampal neurons. A-B: Frontal MRI slices showing the sites where the 111 parietal neurons were recorded, in the 2 monkeys (N_monkey K_=57, N_monkey S_=54). 1 pixel corresponds to 1x1mm. C-D: Neural place maps of PPC (E, in red) and HPC (F, in green) neurons whose activity were not modulated by monkey’s position (not significant position information content). The color axis represents the firing rate (maximal firing rate, in spikes/s, indicated on the bottom left of each map). For a better visualization, data are displayed from the 5th to the 99th percentiles. E-F: Same conventions as for C-D, for PPC (E, in red) and HPC (F, in green) position-modulated cells displaying a significantly symmetrical activity across the whole maze (symmetry index, SI, indicated on the bottom right of each map). G-H: Same conventions, as for E-F, for PPC (G, in red) and HPC (H, in green) position-modulated cells displaying a significantly asymmetrical activity in the inbound paths. I-J: Same conventions, for PPC (I, in red) and HPC (J, in green) position-modulated cells displaying a significantly asymmetrical activity in the outbound paths.

**Figure S2.**
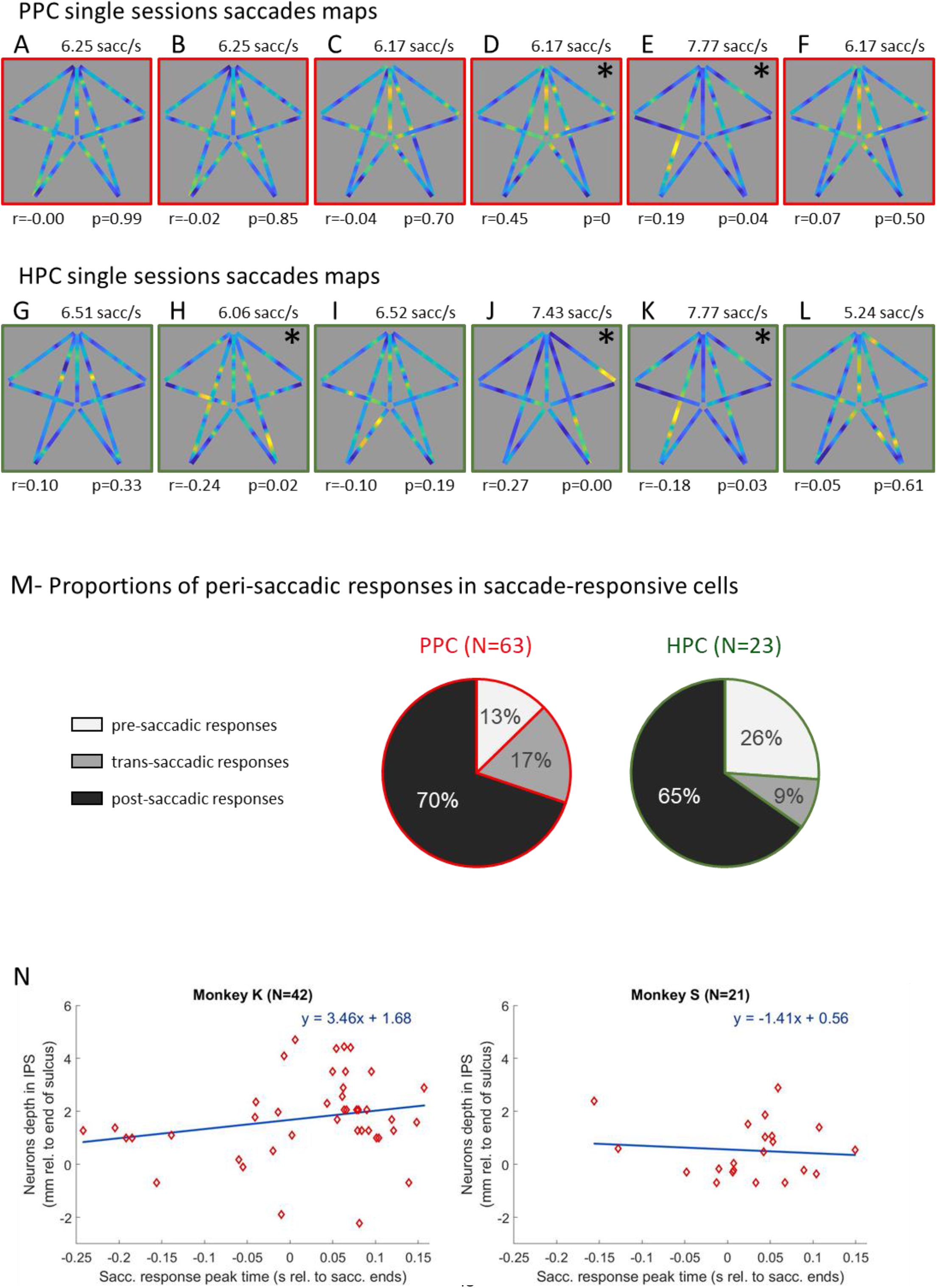
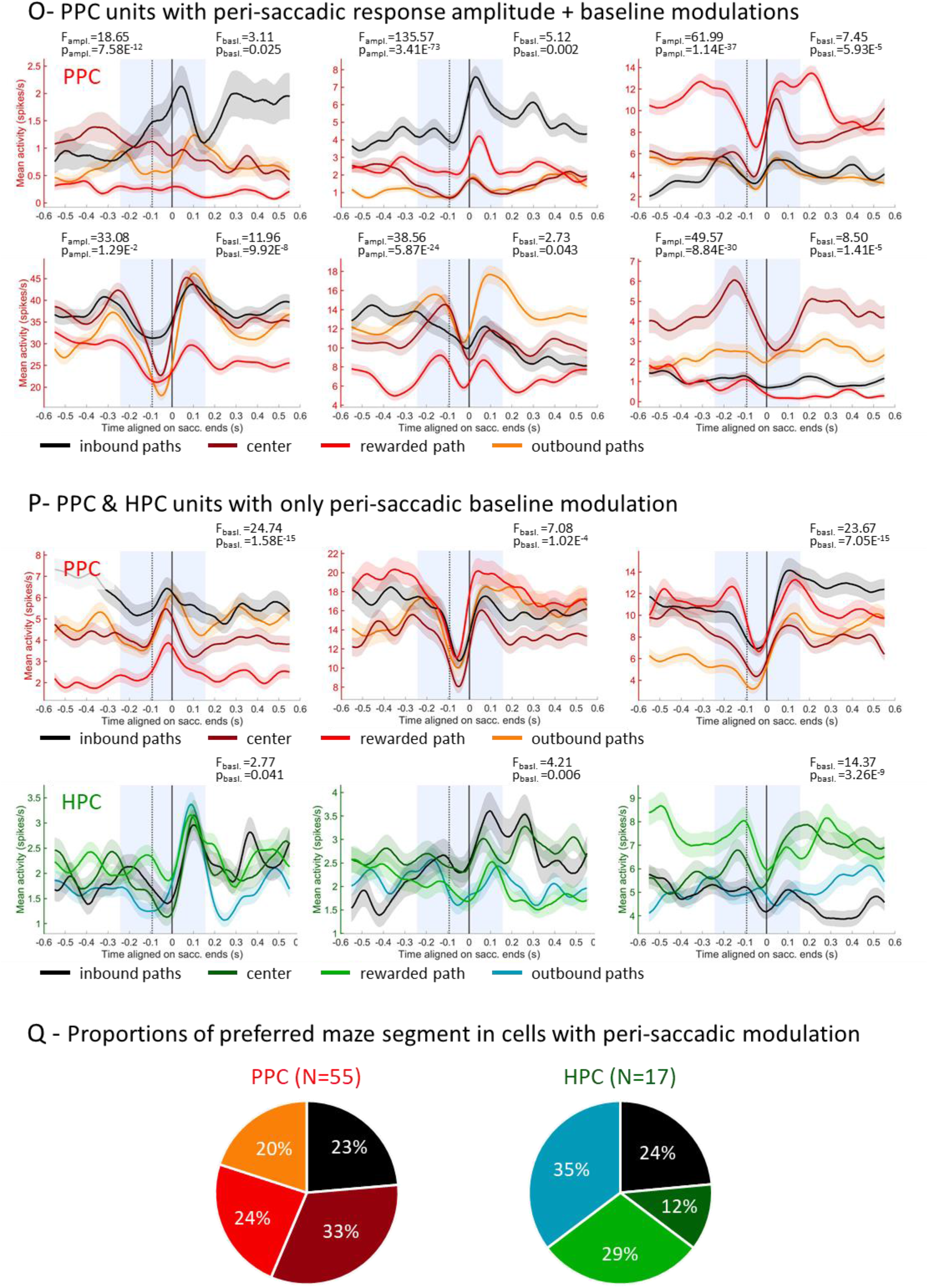
Saccade-related properties of the neurons. A-F: Maps representing the behavioral saccade rate as a function of the animal’s position in the maze for example sessions during which neurons of Figures 1B-G were recorded (i.e. « saccades maps »). The saccade frequency map and the neural place map (Figures 1B-G) were significantly correlated when indicated by the asterisk (permutation tests, p-values on the bottom right of each map, and Person’s correlation coefficient on the bottom left). For a better visualization, data are displayed from the 5th to the 99th percentiles. G-L: Same conventions as for A-F, for HPC example sessions, corresponding to cells of Figures 1H-M. M: Proportions of neurons displaying a pre-(light grey), trans-(medium grey) or post-(black) saccadic peak activity, among the PPC (N=63; in red) and HPC (N=23; in green) saccade-responsive cells. N: Times of the peak activity of the saccade-responsive PPC neurons, relative to saccade ends, as a function of neurons’ anatomical depth in the intraparietal sulcus, relatively to the end of the sulcus, for each monkey (N_monkey K_=42, N_monkey S_=21). The linear regression (in blue) shows no significant effect of the depth on the peak time (Pearson correlation: r_monkey K_=0.22, p_monkey K_=0.16; r_monkey S_=-0.097, p_monkey S_=0.68). O: Averaged activity of example PPC cells, aligned on saccade ends (black solid line). The black dotted line represents the average time of saccades starts (-93ms). The standard errors are indicated in light color. The saccade-related activity is shown for different maze segments (inbound paths in black, center in burgundy, rewarding path in red, outbound paths in orange). All the cells displayed a significant difference of response peak amplitude (1-way-ANOVA, 3df; statistics and p-values indicated on the top left of the figures) as well as a significant difference in peri-saccadic response rate (1-way-ANOVA, 3df; statistics and p-values indicated on the top right of the figures) in the response window (in light blue). P: Same averaged activity as in O, for PPC (top, in reds) and HPC (bottom, in greens) cells that displayed a significant difference in peri-saccadic response rate (1-way-ANOVA, 3df; statistics and p-values indicated on the top right of the figures), but no difference in response peak amplitude. Q: Proportions of maze segment for which the saccade-related activity was the highest for the PPC (N=55) and HPC (N=17) cells displaying a selectivity in peri-saccadic response. In PPC, the distribution was uniform (Chi-squared goodness-of-fit: X²_3df_=1.95, p=0.58). In HPC, the sample size of each category was too small to perform the test.

**Figure S3.**
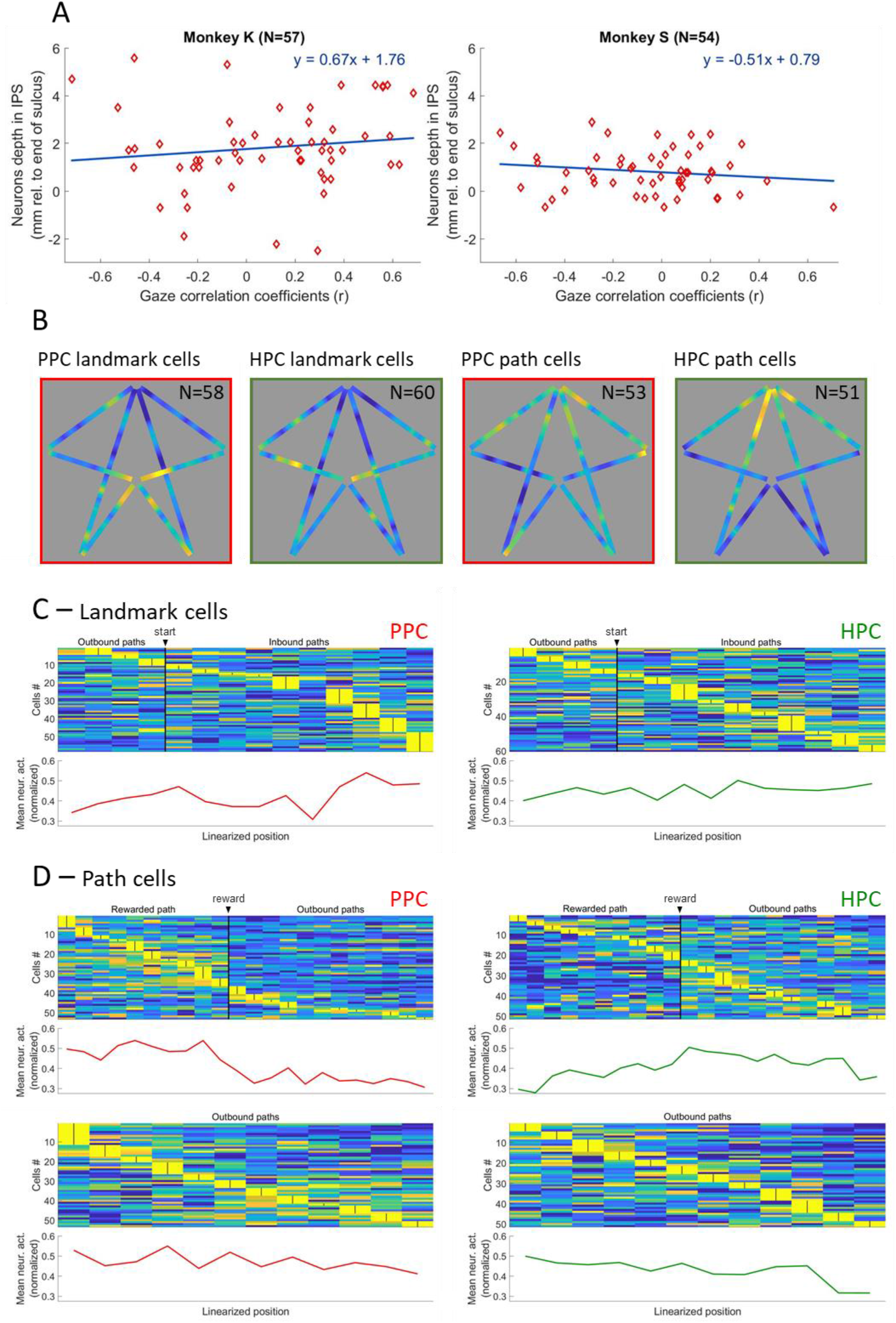
Anatomical, frequency and spatial properties of the landmark and path cells. A: Coefficients of the correlation between neurons’ gaze-related activity and the reference sine-wave, as a function of neurons’ anatomical depth in the intraparietal sulcus, relatively to the end of the sulcus, for each monkey. Positive coefficients correspond to landmark cells, and negative ones to path cells. The linear regression (in blue) shows no significant effect of the depth on the peak time (Pearson correlation: r_monkey K_=0.13, p_monkey K_=0.32; r_monkey S_=-0.16, p_monkey S_=0.25). B: Averaged normalized neural place maps of PPC and HPC landmark and path cells, after removing of the rewarded path from the maps. The two maps of the landmark cells were significantly correlated (Pearson correlation: r=0.28, permutation test: p=0.010), as well as the two maps of the path cells (r=0.20, p=0.038). C: Normalized population maps (top) of the PPC (N=58; left, in red) and HPC (N=60; right, in green) landmark cells, sorted by temporal order of peak activity for a section comprising the end of the outbound paths and the full inbound paths, and the corresponding average of the normalized activity, as a function of monkey’s position (bottom). The distributions of the activity peaks were different between the two regions (Kolmogorov-Smirnov test: D=0.27, p=0.025). D: Same population maps and averaged normalized neuronal activity, for PPC (N=53; left, in red) and HPC (N=51; right, in green) path cells, for positions comprised in the rewarded path and the outbound paths (top), and for the outbound paths only (bottom). The distributions of the activity peaks were different between the two regions when taking in account the rewarded path (D=0.26, p=0.043), but not when excluded (D=0.16, p=0.45).

## References

Acharya L, Aghajan ZM, Vuong C, Moore JJ, Mehta MR. Causal Influence of Visual Cues on Hippocampal Directional Selectivity. Cell 164: 197–207, 2016.

Ammassari-Teule M, Save E, de Marsanich B, Thinus-Blanc C. Posterior parietal cortex lesions severely disrupt spatial learning in DBA mice characterized by a genetic hippocampal dysfunction. Behavioural Brain Research 95: 85–90, 1998.

Arbib MA. From visual affordances in monkey parietal cortex to hippocampo-parietal interactions underlying rat navigation. 1997.

Arleo A, Rondi-Reig L. Multimodal sensory integration and concurrent navigation strategies for spatial cognition in real and artificial organisms. J Integr Neurosci 06: 327–366, 2007.

Avillac M, Denève S, Olivier E, Pouget A, Duhamel J-R. Reference frames for representing visual and tactile locations in parietal cortex. Nature Neuroscience 8: 941–949, 2005.

Barash S, Bracewell RM, Fogassi L, Gnadt JW, Andersen RA. Saccade-related activity in the lateral intraparietal area. I. Temporal properties; comparison with area 7a. Journal of Neurophysiology 66: 1095–1108, 1991.

Baumann O, Mattingley JB. Dissociable roles of the hippocampus and parietal cortex in processing of coordinate and categorical spatial information. Front Cell Neurosci 8, 2014.

Ben Hamed S. Visual Receptive Field Modulation in the Lateral Intraparietal Area during Attentive Fixation and Free Gaze. Cerebral Cortex 12: 234–245, 2002.

Ben Hamed S, Duhamel J-R, Bremmer F, Graf W. Representation of the visual field in the lateral intraparietal area of macaque monkeys: a quantitative receptive field analysis. Exp Brain Res 140: 127–144, 2001.

Bendiksby MS, Platt ML. Neural correlates of reward and attention in macaque area LIP. Neuropsychologia 44: 2411–2420, 2006.

Bicanski A, Burgess N. Neuronal vector coding in spatial cognition. Nat Rev Neurosci 21: 453–470, 2020.

Bremmer F. Navigation in space – the role of the macaque ventral intraparietal area. The Journal of Physiology 566: 29–35, 2005.

Bremmer F, Distler C, Hoffmann K-P. Eye Position Effects in Monkey Cortex. II. Pursuit- and Fixation-Related Activity in Posterior Parietal Areas LIP and 7A. Journal of Neurophysiology 77: 962–977, 1997.

Bremmer F, Duhamel J-R, Hamed SB, Graf W. Heading encoding in the macaque ventral intraparietal area (VIP). European Journal of Neuroscience 16: 1554–1568, 2002a.

Bremmer F, Kaminiarz A, Klingenhoefer S, Churan J. Decoding Target Distance and Saccade Amplitude from Population Activity in the Macaque Lateral Intraparietal Area (LIP) [Online]. Frontiers in Integrative Neuroscience 10, 2016https://www.frontiersin.org/articles/10.3389/fnint.2016.00030 [24 Aug. 2023].

Bremmer F, Klam F, Duhamel J-R, Hamed SB, Graf W. Visual–vestibular interactive responses in the macaque ventral intraparietal area (VIP). European Journal of Neuroscience 16: 1569–1586, 2002b.

Bremmer F, Kubischik M, Hoffmann K-P, Krekelberg B. Neural Dynamics of Saccadic Suppression. J Neurosci 29: 12374–12383, 2009.

Bridgeman B, Hendry D, Stark L. Failure to detect displacement of the visual world during saccadic eye movements. Vision Research 15: 719–722, 1975.

Brincat SL, Miller EK. Frequency-specific hippocampal-prefrontal interactions during associative learning. Nat Neurosci 18: 576–581, 2015.

Burgess N, O’Keefe J, Arbib MA. From visual affordances in monkey parietal cortex to hippocampo–parietal interactions underlying rat navigation. Philosophical Transactions of the Royal Society of London Series B: Biological Sciences 352: 1429–1436, 1997.

Calton JL, Taube JS. Where am I and how will I get there from here? A role for posterior parietal cortex in the integration of spatial information and route planning. Neurobiology of Learning and Memory 91: 186–196, 2009.

Camara E, Rodriguez-Fornells A, Münte T. Functional connectivity of reward processing in the brain [Online]. Frontiers in Human Neuroscience 2, 2009https://www.frontiersin.org/articles/10.3389/neuro.09.019.2008 [2 May 2023].

Chen A, DeAngelis GC, Angelaki DE. Representation of Vestibular and Visual Cues to Self-Motion in Ventral Intraparietal Cortex. Journal of Neuroscience 31: 12036–12052, 2011.

Chen G, King JA, Burgess N, O’Keefe J. How vision and movement combine in the hippocampal place code. Proceedings of the National Academy of Sciences 110: 378–383, 2013.

Cohen YE, Andersen RA. A common reference frame for movement plans in the posterior parietal cortex. Nature Reviews Neuroscience 3: 553–562, 2002.

Colby CL, Duhamel J-R. Heterogeneity of extrastriate visual areas and multiple parietal areas in the Macaque monkey. Neuropsychologia 29: 517–537, 1991.

Corrigan BW, Gulli RA, Doucet G, Mahmoudian B, Abbass M, Roussy M, Luna R, Sachs AJ, Martinez-Trujillo JC. View cells in the hippocampus and prefrontal cortex of macaques during virtual navigation. Hippocampus 33: 573–585, 2023.

Corrigan BW, Gulli RA, Doucet G, Martinez-Trujillo JC. Characterizing eye movement behaviors and kinematics of non-human primates during virtual navigation tasks. Journal of Vision 17: 15, 2017.

Corrigan BW, Gulli RA, Doucet G, Roussy M, Luna R, Pradeepan KS, Sachs AJ, Martinez-Trujillo JC. Distinct neural codes in primate hippocampus and lateral prefrontal cortex during associative learning in virtual environments. Neuron 110: 2155–2169.e4, 2022.

Cushman JD, Aharoni DB, Willers B, Ravassard P, Kees A, Vuong C, Popeney B, Arisaka K, Mehta MR. Multisensory Control of Multimodal Behavior: Do the Legs Know What the Tongue Is Doing? PLoS ONE 8: e80465, 2013.

Doucet G, Gulli RA, Corrigan BW, Duong LR, Martinez-Trujillo JC. Modulation of local field potentials and neuronal activity in primate hippocampus during saccades. Hippocampus 30: 192–209, 2020.

Duhamel, Colby CL, Goldberg ME. The updating of the representation of visual space in parietal cortex by intended eye movements. Science 255: 90–92, 1992.

Duhamel J-R, Bremmer F, Hamed SB, Graf W. Spatial invariance of visual receptive fields in parietal cortex neurons. Nature 389: 845, 1997.

Duhamel J-R, Colby CL, Goldberg ME. Ventral Intraparietal Area of the Macaque: Congruent Visual and Somatic Response Properties. Journal of Neurophysiology 79: 126–136, 1998.

Dupret D, O’Neill J, Pleydell-Bouverie B, Csicsvari J. The reorganization and reactivation of hippocampal maps predict spatial memory performance. Nat Neurosci 13: 995–1002, 2010.

Ekstrom AD. Why vision is important to how we navigate: Human Spatial Navigation and Vision. Hippocampus 25: 731–735, 2015.

Falkner AL, Goldberg ME, Krishna BS. Spatial Representation and Cognitive Modulation of Response Variability in the Lateral Intraparietal Area Priority Map. Journal of Neuroscience 33: 16117–16130, 2013.

Fanini A, Assad JA. Direction Selectivity of Neurons in the Macaque Lateral Intraparietal Area. Journal of Neurophysiology 101: 289–305, 2009.

Goldberg ME, Bisley JW, Powell KD, Gottlieb J. Chapter 10 Saccades, salience and attention: the role of the lateral intraparietal area in visual behavior. In: Progress in Brain Research. Elsevier, p. 157–175.

Gottlieb J, Hayhoe M, Hikosaka O, Rangel A. Attention, Reward, and Information Seeking. J Neurosci 34: 15497–15504, 2014.

Gottlieb JP, Kusunoki M, Goldberg ME. The representation of visual salience in monkey parietal cortex. Nature 391: 481–484, 1998.

Grefkes C, Fink GR. The functional organization of the intraparietal sulcus in humans and monkeys. J Anat 207: 3–17, 2005.

Gulli RA, Duong LR, Corrigan BW, Doucet G, Williams S, Fusi S, Martinez-Trujillo JC. Context-dependent representations of objects and space in the primate hippocampus during virtual navigation. Nat Neurosci 23: 103–112, 2020.

Hagan MA, Dean HL, Pesaran B. Spike-field activity in parietal area LIP during coordinated reach and saccade movements. J Neurophysiol 107: 1275–1290, 2012.

Harvey CD, Coen P, Tank DW. Choice-specific sequences in parietal cortex during a virtual-navigation decision task. Nature 484: 62–68, 2012.

Hayhoe M, Ballard D. Eye movements in natural behavior. Trends in Cognitive Sciences 9: 188–194, 2005.

Heiser LM, Colby CL. Spatial Updating in Area LIP Is Independent of Saccade Direction. Journal of Neurophysiology 95: 2751–2767, 2006.

Henderson JM, Williams CC, Falk RJ. Eye movements are functional during face learning. Mem Cogn 33: 98–106, 2005.

Herrington TM, Masse NY, Hachmeh KJ, Smith JET, Assad JA, Cook EP. The Effect of Microsaccades on the Correlation between Neural Activity and Behavior in Middle Temporal, Ventral Intraparietal, and Lateral Intraparietal Areas. J Neurosci 29: 5793–5805, 2009.

Hoffman KL. Saccades during visual exploration align hippocampal 3-8Hz rhythms in human and non-human primates. Frontiers in Systems Neuroscience 10, 2013.

Hok V, Lenck-Santini P-P, Roux S, Save E, Muller RU, Poucet B. Goal-Related Activity in Hippocampal Place Cells. J Neurosci 27: 472–482, 2007.

Hollup SA, Molden S, Donnett JG, Moser M-B, Moser EI. Accumulation of Hippocampal Place Fields at the Goal Location in an Annular Watermaze Task. J Neurosci 21: 1635–1644, 2001.

Hori E, Nishio Y, Kazui K, Umeno K, Tabuchi E, Sasaki K, Endo S, Ono T, Nishijo H. Place-related neural responses in the monkey hippocampal formation in a virtual space. Hippocampus 15: 991–996, 2005.

Hurley N, Rickard S. Comparing Measures of Sparsity. IEEE Transactions on Information Theory 55: 4723–4741, 2009.

Ibbotson MR, Crowder NA, Cloherty SL, Price NSC, Mustari MJ. Saccadic Modulation of Neural Responses: Possible Roles in Saccadic Suppression, Enhancement, and Time Compression. J Neurosci 28: 10952–10960, 2008.

Insausti R, Amaral DG, Cowan WM. The entorhinal cortex of the monkey: II. Cortical afferents. Journal of Comparative Neurology 264: 356–395, 1987.

Ito J, Maldonado P, Singer W, Grün S. Saccade-Related Modulations of Neuronal Excitability Support Synchrony of Visually Elicited Spikes. Cerebral Cortex 21: 2482–2497, 2011.

Jutras MJ, Fries P, Buffalo EA. Oscillatory activity in the monkey hippocampus during visual exploration and memory formation. Proceedings of the National Academy of Sciences 110: 13144–13149, 2013.

Katz CN, Schjetnan AGP, Patel K, Barkley V, Hoffman KL, Kalia SK, Duncan KD, Valiante TA. A corollary discharge mediates saccade-related inhibition of single units in mnemonic structures of the human brain. Current Biology 32: 3082–3094.e4, 2022.

Knudsen EB, Wallis JD. Hippocampal neurons construct a map of an abstract value space. Cell 184: 4640–4650.e10, 2021.

Kobayashi Y, Amaral DG. Macaque monkey retrosplenial cortex: II. Cortical afferents. Journal of Comparative Neurology 466: 48–79, 2003.

Kobayashi Y, Amaral DG. Macaque monkey retrosplenial cortex: III. Cortical efferents. Journal of Comparative Neurology 502: 810–833, 2007.

Kravitz DJ, Saleem KS, Baker CI, Mishkin M. A new neural framework for visuospatial processing. Nature Reviews Neuroscience 12: 217–230, 2011.

Krumin M, Lee JJ, Harris KD, Carandini M. Decision and navigation in mouse parietal cortex. Elife 7, 2018.

Kusunoki M, Goldberg ME. The Time Course of Perisaccadic Receptive Field Shifts in the Lateral Intraparietal Area of the Monkey. Journal of Neurophysiology 89: 1519–1527, 2003.

Lakshminarasimhan KJ, Avila E, Neyhart E, DeAngelis GC, Pitkow X, Angelaki DE. Tracking the Mind’s Eye: Primate Gaze Behavior during Virtual Visuomotor Navigation Reflects Belief Dynamics. Neuron 106: 662–674.e5, 2020.

Leathers ML, Olson CR. In monkeys making value-based decisions, LIP neurons encode cue salience and not action value. Science 338: 132–135, 2012.

Lee JJ, Krumin M, Harris KD, Carandini M. Task specificity in mouse parietal cortex. Neuron S0896627322006626, 2022.

Li L, Warren WH. Perception of heading during rotation: sufficiency of dense motion parallax and reference objects. Vision Research 40: 3873–3894, 2000.

Liu Z-X, Shen K, Olsen RK, Ryan JD. Visual Sampling Predicts Hippocampal Activity. J Neurosci 37: 599–609, 2017.

Loftus GR. Eye fixations and recognition memory for pictures. Cognitive Psychology 3: 525– 551, 1972.

Mao D, Avila E, Caziot B, Laurens J, Dickman JD, Angelaki DE. Spatial modulation of hippocampal activity in freely moving macaques. Neuron 109: 3521–3534.e6, 2021.

Markus EJ, Qin YL, Leonard B, Skaggs WE, McNaughton BL, Barnes CA. Interactions between location and task affect the spatial and directional firing of hippocampal neurons. J Neurosci 15: 7079–7094, 1995.

Martinez-Trujillo J. Corollary discharge: Linking saccades and memory circuits in the human brain. Current Biology 32: R774–R776, 2022.

McNaughton BL, Mizumori SJY, Barnes CA, Leonard BJ, Marquis M, Green EJ. Cortical Representation of Motion during Unrestrained Spatial Navigation in the Rat. Cerebral Cortex 4: 27–39, 1994.

Meister MLR, Buffalo EA. Getting directions from the hippocampus: The neural connection between looking and memory. Neurobiology of Learning and Memory 134: 135–144, 2016.

Melcher D. Persistence of visual memory for scenes. Nature 412: 401–401, 2001.

Mullette-Gillman OA, Cohen YE, Groh JM. Eye-Centered, Head-Centered, and Complex Coding of Visual and Auditory Targets in the Intraparietal Sulcus. Journal of Neurophysiology 94: 2331–2352, 2005.

Munuera J, Duhamel J-R. The role of the posterior parietal cortex in saccadic error processing. Brain Struct Funct 225: 763–784, 2020.

Nitz DA. Tracking Route Progression in the Posterior Parietal Cortex. Neuron 49: 747–756, 2006.

O’Keefe J, Burgess N. Geometric determinants of the place fields of hippocampal neurons. Nature 381: 425–428, 1996.

O’Keefe J, Dostrovsky J. The hippocampus as a spatial map. Preliminary evidence from unit activity in the freely-moving rat. Brain Research 34: 171–175, 1971.

O’Keefe J, Nadel L. Précis of O’Keefe & Nadel’s The hippocampus as a cognitive map. Behavioral and Brain Sciences 2: 487–494, 1979.

Olsen RK, Sebanayagam V, Lee Y, Moscovitch M, Grady CL, Rosenbaum RS, Ryan JD. The relationship between eye movements and subsequent recognition: Evidence from individual differences and amnesia. Cortex 85: 182–193, 2016.

Pertzov Y, Avidan G, Zohary E. Accumulation of visual information across multiple fixations. Journal of Vision 9: 2–2, 2009.

Platt ML, Glimcher PW. Neural correlates of decision variables in parietal cortex. Nature 400: 233–238, 1999.

Rajkai C, Lakatos P, Chen C-M, Pincze Z, Karmos G, Schroeder CE. Transient Cortical Excitation at the Onset of Visual Fixation. Cerebral Cortex 18: 200–209, 2008.

Ravassard P, Kees A, Willers B, Ho D, Aharoni DA, Cushman J, Aghajan ZM, Mehta MR. Multisensory control of hippocampal spatiotemporal selectivity. Science 340: 1342–1346, 2013.

Ringo JL, Sobotka S, Diltz MD, Bunce CM. Eye movements modulate activity in hippocampal, parahippocampal, and inferotemporal neurons. J Neurophysiol 71: 1285–1288, 1994.

Rockland KS, Van Hoesen GW. Some Temporal and Parietal Cortical Connections Converge in CA1 of the Primate Hippocampus. Cereb Cortex 9: 232–237, 1999.

Rogers JL, Kesner RP. Hippocampal–parietal cortex interactions: Evidence from a disconnection study in the rat. Behavioural Brain Research 179: 19–27, 2007.

Rucci M, Poletti M. Control and Functions of Fixational Eye Movements. Annu Rev Vis Sci 1: 499–518, 2015.

Ryals AJ, Wang JX, Polnaszek KL, Voss JL. Hippocampal contribution to implicit configuration memory expressed via eye movements during scene exploration. Hippocampus 25: 1028–1041, 2015.

Sato N, Sakata H, Tanaka Y, Taira M. Navigation in virtual environment by the macaque monkey. Behavioural Brain Research 153: 287–291, 2004.

Save E, Paz-Villagran V, Alexinsky T, Poucet B. Functional interaction between the associative parietal cortex and hippocampal place cell firing in the rat. European Journal of Neuroscience 21: 522–530, 2005.

Save E, Poucet B. Involvement of the hippocampus and associative parietal cortex in the use of proximal and distal landmarks for navigation. Behavioural Brain Research 109: 195–206, 2000.

Selemon LD, Goldman-Rakic PS. Common cortical and subcortical targets of the dorsolateral prefrontal and posterior parietal cortices in the rhesus monkey: evidence for a distributed neural network subserving spatially guided behavior. J Neurosci 8: 4049–4068, 1988.

Sereno MI, Pitzalis S, Martinez A. Mapping of Contralateral Space in Retinotopic Coordinates by a Parietal Cortical Area in Humans. Science 294: 1350–1354, 2001.

Shelley LE, Barr CI, Nitz DA. Cortical and hippocampal dynamics under logical fragmentation of environmental space. Neurobiology of Learning and Memory 189: 107597, 2022.

Shelley LE, Nitz DA. Locomotor action sequences impact the scale of representation in hippocampus and posterior parietal cortex. Hippocampus 31: 677–689, 2021.

Skaggs W, McNaughton B, Gothard K. An Information-Theoretic Approach to Deciphering the Hippocampal Code [Online]. In: Advances in Neural Information Processing Systems. Morgan-Kaufmannhttps://proceedings.neurips.cc/paper_files/paper/1992/hash/5dd9db5e033da9c6fb5ba83c7a7ebea9-Abstract.html [6 Nov. 2023].

Sobotka S, Nowicka A, Ringo JL. Activity linked to externally cued saccades in single units recorded from hippocampal, parahippocampal, and inferotemporal areas of macaques. J Neurophysiol 78: 2156–2163, 1997.

Stachenfeld KL, Botvinick MM, Gershman SJ. The hippocampus as a predictive map. Nat Neurosci 20: 1643–1653, 2017.

Steenrod SC, Phillips MH, Goldberg ME. The lateral intraparietal area codes the location of saccade targets and not the dimension of the saccades that will be made to acquire them. Journal of Neurophysiology 109: 2596–2605, 2013.

Sunkara A, DeAngelis GC, Angelaki DE. Joint representation of translational and rotational components of optic flow in parietal cortex. Proceedings of the National Academy of Sciences 113: 5077–5082, 2016.

Taillade M, N’Kaoua B, Gross C. Navigation strategy in macaque monkeys: An exploratory experiment in virtual reality. Journal of Neuroscience Methods 326: 108336, 2019.

Thinus-Blanc C, Save E, Rossi-Arnaud C, Tozzi A, Ammassari-Teule M. The differences shown by C57BL/6 and DBA/2 inbred mice in detecting spatial novelty are subserved by a different hippocampal and parietal cortex interplay. Behavioural Brain Research 80: 33–40, 1996.

Van Hoesen GW. The parahippocampal gyrus: New observations regarding its cortical connections in the monkey. Trends in Neurosciences 5: 345–350, 1982.

Wardak C, Olivier E, Duhamel J-R. Saccadic Target Selection Deficits after Lateral Intraparietal Area Inactivation in Monkeys. J Neurosci 22: 9877–9884, 2002.

Wilber AA, Clark BJ, Forster TC, Tatsuno M, McNaughton BL. Interaction of Egocentric and World-Centered Reference Frames in the Rat Posterior Parietal Cortex. J Neurosci 34: 5431–5446, 2014.

Wirth S, Avsar E, Chiu CC, Sharma V, Smith AC, Brown E, Suzuki WA. Trial Outcome and Associative Learning Signals in the Monkey Hippocampus. Neuron 61: 930–940, 2009.

Wirth S, Baraduc P, Planté A, Pinède S, Duhamel J-R. Gaze-informed, task-situated representation of space in primate hippocampus during virtual navigation. PLOS Biology 15: e2001045, 2017.

Wittmann BC, Schott BH, Guderian S, Frey JU, Heinze H-J, Düzel E. Reward-Related fMRI Activation of Dopaminergic Midbrain Is Associated with Enhanced Hippocampus-Dependent Long-Term Memory Formation. Neuron 45: 459–467, 2005.

Wolbers T, Hegarty M, Büchel C, Loomis JM. Spatial updating: how the brain keeps track of changing object locations during observer motion. Nature Neuroscience 11: 1223–1230, 2008.

Zanos TP, Mineault PJ, Nasiotis KT, Guitton D, Pack CC. A Sensorimotor Role for Traveling Waves in Primate Visual Cortex. Neuron 85: 615–627, 2015.

Zhang M, Wang X, Goldberg ME. A spatially nonselective baseline signal in parietal cortex reflects the probability of a monkey’s success on the current trial. Proceedings of the National Academy of Sciences 111: 8967–8972, 2014.

Zhou Y, Liu Y, Wu S, Zhang M. Neuronal Representation of the Saccadic Timing Signals in Macaque Lateral Intraparietal Area. Cereb Cortex 28: 2887–2900, 2018.

Zhu S, Lakshminarasimhan KJ, Arfaei N, Angelaki DE. Eye movements reveal spatiotemporal dynamics of visually-informed planning in navigation. eLife 11: e73097, 2022.

Zhu SL, Lakshminarasimhan KJ, Angelaki DE. Computational cross-species views of the hippocampal formation. Hippocampus n/a, 2023.

